# Tests for Segregation Distortion in Tetraploid F1 Populations

**DOI:** 10.1101/2024.02.07.579361

**Authors:** David Gerard, Mira Thakkar, Luis Felipe V. Ferrão

## Abstract

Genotype data from tetraploid F1 populations are often collected in breeding programs for mapping and genomic selection purposes. A common quality control procedure in these groups is to compare empirical genotype frequencies against those predicted by Mendelian segregation, where SNPs detected to have “segregation distortion” are discarded. However, current tests for segregation distortion are insufficient in that they do not account for double reduction and preferential pairing, two meiotic processes in polyploids that naturally change gamete frequencies, leading these tests to detect segregation distortion too often. Current tests also do not account for genotype uncertainty, again leading these tests to detect segregation distortion too often. Here, we incorporate double reduction, preferential pairing, and genotype uncertainty in likelihood ratio and Bayesian tests for segregation distortion. Our methods are implemented in a user-friendly R package, menbayes. We demonstrate the superiority of our methods to those currently used in the literature on both simulations and real data.

## 1 Introduction

Polyploids, organisms containing more than two sets of chromosomes, play a dominant role in many sectors of agriculture [Udall and Wendel, 2006]. Consequently, numerous breeding programs are dedicated to the agricultural improvement of polyploids [Ferrão et al., 2018, Shirasawa et al., 2017, Amadeu et al., 2021, Lau et al., 2022]. In these programs, breeders frequently generate “F1 populations” of full siblings for various tasks, such as QTL mapping [Amadeu et al., 2021], linkage mapping [Bourke et al., 2018, Mollinari and Garcia, 2019], and genomic selection [Ferrão et al., 2021], all of which are crucial for crop improvement.

In these F1 populations, offspring genotypes should roughly adhere to the laws of Mendelian segregation [Mendel, 1866]. Hence, it is customary to use a chi-squared test to compare observed offspring genotype frequencies with those predicted by Mendelian segregation to identify problematic SNPs caused, for example, by sequencing errors, mapping biases, or amplification biases [Bourke et al., 2015, Cappai et al., 2020, Mollinari et al., 2020, Batista et al., 2021, e.g.]. Such deviations are referred to as “segregation distortion.” However, there are two significant limitations to using the chi-squared test in these scenarios. First, many polyploids naturally undergo double reduction and (partial) preferential pairing [Voorrips and Maliepaard, 2012], two meiotic processes that can lead to deviations from classical gamete frequencies even for well-behaved SNPs. The resulting offspring genotype frequencies heavily depend on the type of polyploid (allo, auto, or segmental) [Doyle and Egan, 2010], necessitating tests for F1 populations that can adapt to these varying types. Second, the chi-squared test does not account for genotype uncertainty, a major concern in polyploid genetics [Gerard et al., 2018, Gerard and Ferrão, 2019] that can adversely impact many genomics methods.

In this paper, we develop a model for the genotype frequencies of a biallelic locus in an F1 tetraploid population that allows for arbitrary levels of double reduction and preferential pairing (Section 2.1). This fills a gap in the literature, as most approaches only account for either double reduction or preferential pairing, but not both (Appendix S1). We harness this new model to develop likelihood ratio tests (LRTs) for segregation distortion, optionally accounting for genotype uncertainty through genotype likelihoods [Li, 2011] (Section 2.2). To take advantage of the benefits of a Bayesian paradigm approach, we further develop Bayesian tests for segregation distortion (Section 2.3). We demonstrate our methods both on simulations (Sections 3.1 and 3.2) and on a dataset of tetraploid blueberries (Section 3.3).

### 1.1 Related work

In the context of modeling preferential pairing and double reduction, previous studies have primarily focused on estimation rather than testing. A comprehensive review of these studies is provided in Appendix S1. In this section, we focus on the related work that emphasizes testing.

Tests have been created to evaluate the related hypothesis of random mating. A likelihood ratio test for random mating was created in Appendix C of Gerard [2022], exact tests were explored in Matoka Nana [2023], and Bayesian tests were developed in Gerard [2023]. Many of the approaches in those papers account for genotype uncertainty. Random mating is applicable to S1 populations (a generation of selfing) as all individuals have their gametes drawn from the same distribution and are randomly selected during fertilization. However, F1 populations violate the random mating hypothesis at loci where parental genotypes differ since the gametes from each parent are drawn from different distributions. Thus, these tests are not generally applicable in our scenario of F1 populations.

The work most closely related to ours, particularly in terms of testing, is likely the tests implemented by the `polymapR` software [Bourke et al., 2018]. This software offers tests for segregation distortion in tetraploids within its function `checkF1(`). The process involves analyzing each segregation pattern, which can be (i) polysomic in both parents, (ii) disomic in both parents, or (iii) polysomic in one and disomic in the other, followed by conducting a chi-squared test based on that specific segregation pattern. This test is performed using only the possible genotypes. For instance, if the potential offspring genotypes from parent genotypes are 0 and 1, but some offspring genotypes of 2 are observed, these genotypes are excluded from the chi-squared test. A separate 1-sided binomial test is conducted for “invalid” genotypes (considering the parental genotypes and their segregation patterns), with an expected proportion of invalid genotypes hard-coded at less than 3%. The product of the *p*-values from both the chi-squared and binomial tests is then calculated, and the maximum of these is used as the indicator of segregation distortion. This method resembles a minimum chi-squared test [Berkson, 1980] where the authors explore the discrete parameter space of fully disomic and fully polysomic parents, albeit using a somewhat ad-hoc criterion. Our approach, in contrast, is more principled and allows for a full exploration of the parameter space of gamete frequencies resulting from both double reduction and partial preferential pairing, rather than limiting to completely polysomic or completely disomic inheritance.

Bourke et al. [2018] also account for genotype uncertainty by using posterior probabilities as inputs, but do so in an ad-hoc way. They sum the posterior probability of each genotype over the individuals to get a total count for each genotype, they then round counts below some pre-set threshold down to zero, and re-normalize the resulting count vector to sum to the sample size of the offspring. They then apply the same approach as in the known genotype case to this estimated vector of counts.

We will show in the Section 3.1 that our approach has advantages to that of Bourke et al. [2018].

## 2 Materials and methods

### 2.1 Generalized gamete frequencies

We begin by describing the hypothesis of no segregation distortion. We assume that we are working with a single biallelic locus, and we are concerned with the genotype frequencies of an F1 population of polyploids at this locus. Let ***q*** = (*q*_0_, *q*_1_, …, *q*_*K*_) be the genotype frequencies of a *K*-ploid F1 population, where *q*_*k*_ is the proportion of offspring expected to have genotype *k*. Each parent provides a gamete to each offspring, and the “gamete frequencies” of parent *j* ∈ {1, 2} will be denoted by ***p***_*j*_ = (*p*_*j*0_, *p*_*j*1_, …, *p*_*j,K/*2_). That is, *p*_*jk*_ is the proportion of parent *j*’s gametes expected to have genotype *k*. Because each offspring genotype is the sum of the two (independent) parental gamete genotypes, we can write ***q*** as a discrete linear convolution of ***p***_1_ and ***p***_2_,

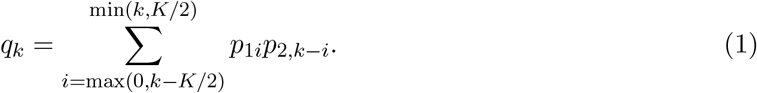

In tetraploids, the subject of our paper, this corresponds to

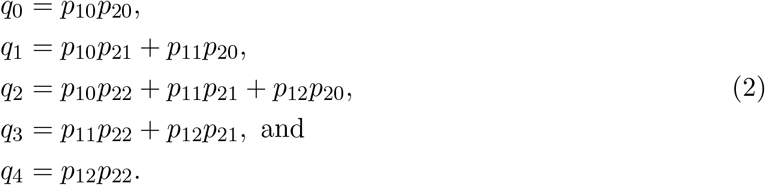

Not all values of ***p***_*j*_ are possible, and models for segregation correspond to models for the ***p***_*j*_ ‘s based on each parental genotype. Let *ℓ*_*j*_ ∈ {0, 1, …, *K*} be the genotype for parent *j*. For true autopolyploids that exhibit strict bivalent pairing, the *p*_*jk*_’s are hypergeometric probabilities [Muller, 1914, Serang et al., 2012],

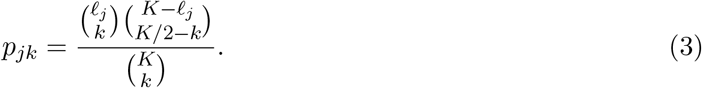

However, polyploids often exhibit some quadrivalent pairing, which can lead to the meiotic process of “double reduction”, the co-migration of sister chromatids segments into the same gamete [Mather, 1935, Stift et al., 2010]. Double reduction alters the gamete frequencies for polyploids. The characterization of these gamete frequencies was described in Fisher and Mather [1943] for autotetraploids and autohexaploids, before being generalized to arbitrary ploidy levels in Huang et al. [2019].

Additionally, many polyploids exhibit partial (or full) preferential pairing, where homologues preferentially (or exclusively) form bivalents during meiosis. Those that exhibit full disomic inheritance are called “allopolyploids” [Doyle and Egan, 2010, Parisod et al., 2010], while those that exhibit partial preferential pairing are called “segmental allopolyploids” [Stebbins, 1947] among other terms [Bourke et al., 2017]. No model yet exists to incorporate both double reduction and preferential pairing at biallelic loci, though Stift et al. [2008] produced a model that incorporates both of these processes in tetraploids when each chromosome is distinguishable.

For one of our contributions, in Appendix S2 we developed a model that incorporates both double reduction and preferential pairing in the gamete frequencies of tetraploids. These frequencies are tabulated in Table 1 in terms of three parameters: the probability of quadrivalent formation, *τ*, the probability of double reduction given quadrivalent formation, *β*, and the probability that chromosomes with the same alleles will pair given bivalent formation, *γ*. We further show that this three parameter model can be reduced to a model with two parameters (Table 2): the double reduction rate, *α*, and the preferential pairing parameter, *ξ*, where a value of *ξ* = 1*/*3 indicates strict polysomic inheritance and values of *ξ* = 0 or 1 indicate strict disomic inheritance. Our model is nicely connected with others in the literature. Our model is derived from that of Stift et al. [2008], reduced to biallelic loci, when the parameters of that model are reinterpreted. Furthermore, when *ξ* = 1*/*3 this model reduces to that of Fisher and Mather [1943] (Appendix S3).

**Table 1:**
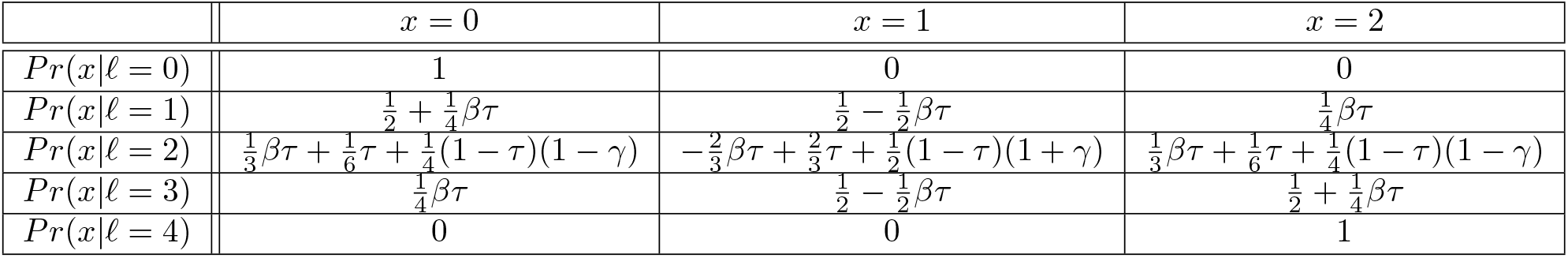
At a single locus for a tetraploid, the distribution of the number, *x*, of alternative alleles sent to an offspring by a parent with dosage *ℓ* = 0, 1, 2, 3, or 4. The probability of quadrivalent formation is *τ, β* is the probability of double reduction given quadrivalent formation, and *γ* is the probability that chromosome pairing occurs along shared alleles given bivalent formation.

**Table 2:** At a single locus for a tetraploid, the distribution of the number, *x*, of alternative alleles sent to an offspring by a parent with dosage *ℓ* = 0, 1, 2, 3, or 4. The double reduction rate is *α* and the preferential pairing parameter is *ξ*. No preferential pairing corresponds to *ξ* = 1*/*3.

| | $x = 0$ | $x = 1$ | $x = 2$ |
| --- | --- | --- | --- |
| $Pr(x\vert \ell = 0)$ | 1 | 0 | 0 |
| $Pr(x\vert \ell = 1)$ | $\frac{1}{2} + \frac{1}{4}\alpha$ | $\frac{1}{2} - \frac{1}{2}\alpha$ | $\frac{1}{4}\alpha$ |
| $Pr(x\vert \ell = 2)$ | $\frac{1}{2}\alpha + \frac{1}{4}(1 - \alpha)(1 - \xi)$ | $\frac{1}{2}(1 - \alpha)(1 + \xi)$ | $\frac{1}{2}\alpha + \frac{1}{4}(1 - \alpha)(1 - \xi)$ |
| $Pr(x\vert \ell = 3)$ | $\frac{1}{4}\alpha$ | $\frac{1}{2} - \frac{1}{2}\alpha$ | $\frac{1}{2} + \frac{1}{4}\alpha$ |
| $Pr(x\vert \ell = 4)$ | 0 | 0 | 1 |

The benefit of our model is that it can account for a wider range of possible gamete frequencies than models that incorporate double reduction alone. That is, some well-behaved SNPs (with some amount of preferential pairing) cannot have their genotype frequencies modeled appropriately with double reduction alone. To see this, consider that for *ℓ* = 2, *p*_0_ = *p*_2_, we can order the gamete frequencies by their value of *p*_1_ = 1 − *p*_0_ − *p*_2_. When accounting for double reduction alone, the range of gamete frequencies when *ℓ* = 2 goes from ***p*** = (2, 5, 2)*/*9 ≈ (0.22, 0.56, 0.22) (for *α* = 1*/*6) to ***p*** = (1, 4, 1)*/*6 ≈ (0.17, 0.67, 0.17) (for *α* = 0). When accounting for both double reduction and preferential pairing, the range of gamete frequencies goes from ***p*** = (1, 2, 1)*/*4 (for *ξ* = 0 and *α* = 0) to ***p*** = (0, 1, 0) (for *ξ* = 1 and *α* = 0). Thus, values of 2*/*3 *< p*_1_ ≤ 1 (and so 0 ≤ *p*_0_, *p*_2_ *<* 1*/*6) and 1*/*2 *< p*_1_ *<* 5*/*9 cannot be accounted for by double reduction alone, but can be accounted for when including preferential pairing.

In Sections 2.2 and 2.3, we will use our new model to construct tests for segregation distortion in F1 populations of tetraploids. There, we will assume that the two parents share a common double reduction rate (*α*), but each has their own preferential pairing parameter (*ξ*_1_ and *ξ*_2_). It would be incorrect to fix *ξ*_1_ to equal *ξ*_2_ due to the interpretation of this parameter in term of pairing frequencies based on allele compositions (see Section 4).

### 2.2 Likelihood ratio tests for segregation distortion

Our goal in this section is to construct LRTs to compare the following two hypotheses.

- *H*_0_: ***p***_*j*_ is defined by Table 2 via parameters *α* and *ξ*_*j*_, and ***q*** is defined by (2).
- *H*_*A*_: Not *H*_0_

When considering *H*_0_, we will denote the functional dependence of ***q*** on *α, ξ*_1_, *ξ*_2_, *ℓ*_1_, and *ℓ*_2_ by ***q***(*α, ξ*_1_, *ξ*_2_, *ℓ*_1_, *ℓ*_2_) if using the two parameter model (Table 2). If using the three parameter model (Table 1), we will denote this dependence by ***q***(*τ, β, γ*_1_, *γ*_2_, *ℓ*_1_, *ℓ*_2_). We construct these tests in three scenarios: one where the genotypes are known, one where parental genotypes are known but offspring genotype uncertainty is represented through genotype likelihoods [Li, 2011], and one where all individuals have genotype uncertainty represented through genotype likelihoods.

We begin with the case when the genotypes are known. Let *x*_*k*_ be the number of individuals with genotype *k* ∈ {0, 1, …, *K*}, which we collect into the vector ***x*** = (*x*_0_, *x*_1_, …, *x*_*K*_). We denote the sample size by 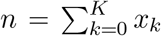. Then, given genotype frequencies ***q***, we have that ***x*** follows a multinomial distribution,

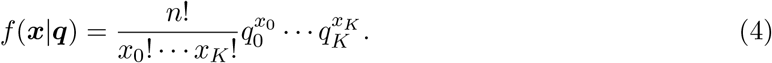

The maximum likelihood estimate of ***q*** under the alternative is 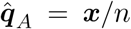. We maximize the likelihood function, *f* (***x***|***q***(*τ, β, γ*_1_, *γ*_2_, *ℓ*_1_, *ℓ*_2_)), over 0 ≤ *τ, γ*_1_, *γ*_2_ ≤ 1 and 0 ≤ *β* ≤ *c*, where *c* is the maximum rate of double reduction. By default, we set *c* = 1*/*6, the maximum under the complete equational segregation model [Mather, 1935]. We do this maximization using gradient ascent to obtain 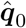. We then obtain the likelihood ratio statistic

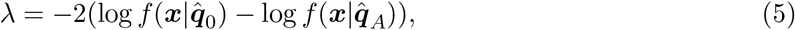

and compare *⋋* to an appropriate *χ* ^2^ distribution to obtain a *p*-value.

Calculating the null distribution of this test is rather difficult, as the parameters under the null might lie on or near the boundary of the parameter space, which requires special considerations [Self and Liang, 1987, Mitchell et al., 2019, Leung and Sturma, 2024]. Thus, we applied the datadependent degrees of freedom strategy of Susko [2013], which we describe now. If *ℓ* _1_, *ℓ* _2_ ∈ {0, 4}, then the number of parameters under the null is 0. If *ℓ* ∈ {1, 2, 3}, then the number of parameters under the null is 1 if the parameters are estimated in the interior of the parameter space (it is never 2 because of the non-identifiability of the model, see Section 4), and is 0 if they are estimated on the boundary of the parameter space. To calculate the number of parameters under the alternative, we note that, if the null were true, some offspring genotypes would be impossible. The test returns a *p*-value of 0 if any of these “impossible” genotypes are observed, otherwise the number of parameters under the alternative is the number of theoretically non-zero elements minus 1. The number of degrees of freedom for the chi-squared test is the difference between the number of parameters under the alternative and under the null. This strategy is guaranteed to asymptotically control Type I error, but might be asymptotically conservative [Susko, 2013].

We now consider the LRT when parental genotypes are known, but offspring use genotype likelihoods [Li, 2011]. Let *g*_*ik*_ be the genotype likelihood for individual *i* = 1, 2, …, *n* for genotype *k* = 0, 1, …, *K*. That is, *g*_*ik*_ is the probability of the data (sequencing, microarray, or otherwise) for individual *i* given that the genotype for that individual is *k*. Then, given these genotype likelihoods, we have the likelihood for these data is

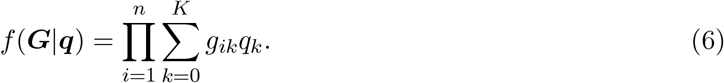

The maximum likelihood estimate of ***q*** under the alternative can be found by the EM algorithm of Li [2011], which we denote by 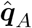. We maximize *f* (***G***|***q***(*τ, β, γ*_1_, *γ*_2_, *ℓ* _1_, *ℓ* _2_)) over 0 ≤ *τ, γ*_1_, *γ*_2_ ≤ 1 and 0 ≤ *β* ≤ *c*, using gradient ascent, to obtain 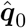. We obtain a *p*-value by comparing the likelihood ratio statistic,

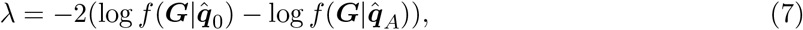

to an appropriate *χ* ^2^ distribution to obtain a *p*-value.

To obtain the number of degrees of freedom of this test, we again take the approach of Susko [2013]. The number of parameters under the null is the same as in the known genotype case. The number of parameters under the alternative is 4 minus the number of the *q*_*k*_’s that are both theoretically 0 under the null and are estimated to be 0 under the alternative. The number of degrees of freedom of the test is the difference between the number of parameters under the alternative and the null. Again, this strategy is guaranteed to asymptotically control from Type I error, but might be asymptotically conservative [Susko, 2013].

We now consider the case when both parents and offspring use genotype likelihoods. Let ***a*** = (*a*_0_, *a*_1_, …, *a*_*K*_) be the genotype likelihoods for parent 1, and let ***b*** = (*b*_0_, *b*_1_, …, *b*_*K*_) be the genotype likelihoods for parent 2. We perform the LRT by first maximizing the following likelihood over the parent genotypes

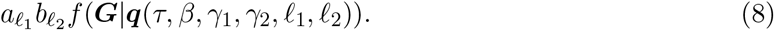

We then run the LRT as if the estimated parent genotypes were the known true parent genotypes.

We further implemented all of these LRTs in the cases when (i) only the double reduction rate (*α*) is known, (ii) only the preferential pairing parameters (*ξ*_1_ and *ξ*_2_) are known, and (iii) both the double reduction rate and the preferential pairing parameters are known.

### 2.3 Bayesian tests for segregation distortion

To take advantage of the many benefits of Bayesian analysis, we developed Bayesian tests for segregation distortion. In particular to our case, Bayesian tests can more easily adapt to non-identifiable models, as this just alters the prior distribution over a parameter space. But there are other advantages, such as ease of interpretability and consistency under the null [O’Hagan, 1994, Section 7.52]. The Bayesian testing paradigm consists of calculating a Bayes Factor (BF) defined as the ratio of marginal likelihoods under the two hypotheses:

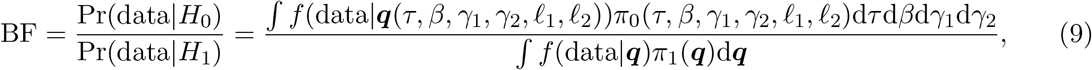

where *π*_0_(*·*) is the prior under the null, *π*_1_(*·*) is the prior under the alternative, and *f* (data|***q***) is one of the likelihoods we consider, either equation (4) or (6). When parent genotypes are not known, we estimate the parent genotypes using maximum likelihood, as in Section 2.2, and use likelihood (6) as if the parent genotypes were known.

Under the alternative, we set the prior over ***q*** to be uniform on the unit 4-simplex. For the null, we need to specify priors over *τ*, the probability of quadrivalent formation, *β*, the probability of double reduction given quadrivalent formation, and *γ*_*j*_, the probability of a `AA:aa` pairing given bivalent formation in parent *j* = 1, 2. Our default selection is as follows,

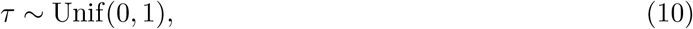

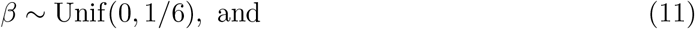

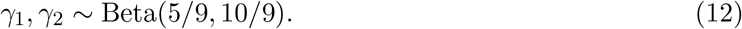

The upper bound on *β* was chosen based on the maximum rate of double reduction, provided by the complete equational segregation model of meiosis [Mather, 1935, Huang et al., 2019]. The prior on the *γ*_*j*_’s was created so that the mean would be 1/3, the value under tetrasomic inheritance (Appendix S3), and so that it would have the same variance as a uniform prior. All of these priors are adjustable by the user if they have additional prior knowledge on the meiotic process they study. E.g., if it is known that only some preferential pairing occurs, then the user could adjust the priors over *γ*_1_ and *γ*_2_ to be more concentrated around 1/3.

Under the alternative, when genotypes are known, the marginal likelihood is the Dirichlet-multinomial [Mosimann, 1962], which can be easily calculated. For all other models and likelihoods, we have to resort to simulation to estimate the marginal likelihoods. We implemented these models, using all three likelihoods and both the null and alternative priors, in Stan [Stan Development Team, 2022a,b]. We estimated marginal likelihoods (and therefore Bayes factors) via bridge sampling [Meng and Wong, 1996, Gronau et al., 2020].

We further implemented all of these Bayesian tests in the cases when (i) only the double reduction rate (*α*) is known, (ii) only the preferential pairing parameters (*ξ*_1_ and *ξ*_2_) are known, and (iii) both the double reduction rate and the preferential pairing parameters are known.

## 3 Results

### 3.1 Null simulations

To evaluate our methods, we ran simulations when the null of no segregation distortion was true. We varied the following parameters:

- The parent genotypes, (*ℓ* _1_, *ℓ* _2_) ∈ {(0, 1), (0, 2), (1, 1), (1, 2), (2, 2)}.
- The sample size, *n* ∈ {20, 200}.
- The double reduction rate, *α* ∈ {0, 1*/*12, 1*/*6}.
- The preferential pairing parameters, 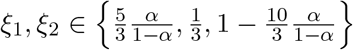. We only varied the preferential pairing parameter *ξ*_*j*_ when *ℓ* _*j*_ = 2. When *α* = 1*/*6, the bounds on the preferential pairing parameter constrains *ξ*_1_ = *ξ*_2_ = 1*/*3 (Theorem S2).
- The read depth, {10, ∞}, where a read depth of ∞ corresponds to the known genotype case.

Each replication, we simulated offspring genotypes using the model of Table 2. When genotypes were not known (a read-depth of 10), we further simulated offspring read-counts using the model of Gerard et al. [2018] under no allele bias, an overdispersion level of 0.01, and a sequencing error rate of 0.01. We then used the method of Gerard et al. [2018] to estimate offspring genotypes and obtain genotype likelihoods. Each replication, we fit the standard chi-squared test for segregation distortion, the `polymapR` test from Section 1.1 [Bourke et al., 2018], our new LRT from Section 2.2, and our new Bayesian test from Section 2.3. For each unique combination of parameter values, we ran 200 replications.

Quantile-quantile plots against the uniform distribution of the *p*-values from the LRT of Section 2.2, the standard chi-squared test, and the `polymapR` test of Section 1.1 [Bourke et al., 2018] are presented in Figures S1–S6. Since the null is true, the *p*-values should lie at or above the *y* = *x* line to control Type I error. Our new LRT is able to control Type I error in all scenarios, often being consistent and only sometimes being conservative (Figures S1–S2). In contrast, the chi-squared test does not control Type I error when there is any double reduction or preferential pairing, and fails to control type I error in almost all scenarios where there is genotype uncertainty (Figures S3–S4). The `polymapR` test fails to control Type I error in some scenarios when genotypes are known, particularly when there is preferential pairing (Figure S5). When genotypes are not known, the `polymapR` test appears to control for Type I error at small samples sizes for many scenarios (likely due to low power), but fails to control for Type I error in most scenarios at larger sample sizes (Figure S6).

Box plots of the log Bayes factors from the Bayesian test of Section 2.3 are presented in Figures S7–S8. Since the null is true, the log Bayes factors should be mostly positive, which is what we see. Furthermore, we see that the log Bayes factors are more positive for larger sample sizes, indicating stronger support for the null hypothesis, which is expected.

### 3.2 Alternative simulations

To evaluate our methods, we ran simulations when the alternative was true. We set the true genotype frequencies at ***q*** = (1, 1, 1, 1, 1)*/*5 and tested for segregation distortion assuming *ℓ* _1_ = *ℓ* _2_ = 2. We varied the sample size *n* ∈ {20, 200} and the read-depth {10, ∞}, where a read depth of ∞ corresponds to the known genotype case. Each replication, we simulated offspring genotypes assuming the appropriate ***q*** from a multinomial distribution. Our procedure for using genotype likelihoods, and the methods we fit each replication, were the same as in Section 3.1. For each unique combination of parameter values, we ran 200 replications.

A table of power at various significance levels for the three methods is presented in Table S1. We see there at the chi-square test is more powerful than the `polymapR` test, which is more powerful than the new LRT. This is the cost of properly controlling for Type I error. However, at larger sample sizes, the power was about 1 for all tests at significance levels at least 0.00001.

Box plots for the log Bayes factors are presented in Figure S9. The Bayes factors are negative in all scenarios, indicating support for the alternative.

### 3.3 Blueberries

We applied our methods on a dataset of F1 tetraploid blueberries (*Vaccinium corymbosum*) (2n = 4x = 48) from Cappai et al. [2020]. The data we considered initially consisted of 21513 SNPs for the *n* = 240 offspring and the 2 parents. We obtained genotype likelihoods using the method of Gerard et al. [2018] with the proportional normal prior [Gerard and Ferrão, 2019]. Markers were were then filtered to remove monomorphic SNPs, defined as those whose maximum genotype frequency was estimated to be greater than 0.95 (20251 remaining SNPs). We then filtered SNPs to keep only loci belonging to the 12 main linkage groups (19524 remaining SNPs). We then ran our LRT (Section 2.2), our Bayesian test (Section 2.3), the standard chi-squared test, and the `polymapR` test for each SNP.

The Bayesian, LRT, and `polymapR` tests generally agree on the amount of segregation distortion in the data. At a Bonferroni adjusted significance level of 0.05, the LRT and `polymapR` indicated a segregation distortion rate of 4.4% and 2.6%, respectively. The Bayesian test had 1.6% of SNPs with a log Bayes factor less than -16 [see Wakefield, 2010, Gerard, 2023, for threshold recommendations for Bayes factors]. In contrast, the chi-squared test using posterior mode genotypes indicated 72.8% of SNPs are in segregation distortion, using a Bonferroni corrected significance level of 0.05.

The likelihood ratio and Bayes tests have more concordance on which SNPs indicate segregation distortion (Figure S10). It is enlightening to see which SNPs `polymapR` and our new methods disagree about. In Figure S11, we provide genotype plots [Gerard et al., 2018] of five SNPs where `polymapR` indicates no segregation distortion while the LRT indicates extreme segregation distortion. In Figure S12 we provide genotype plots of five SNPs where `polymapR` indicates extreme segregation distortion while the LRT indicates no segregation distortion. The *p*-values of the various tests for these SNPs are provided in Table S2.

Generally, since `polymapR` does not account for double reduction, it detects segregation distortion in SNPs that seem to have high rates of double reduction. Examples of these are presented in the last 5 rows of Table S2. These are all simplex *×* nullplex markers that roughly exhibit the 13:10:1 segregation ratios one would expect at a double reduction rate of *α* = 1*/*6, and so our LRT and Bayes test correctly indicate that there is no evidence of segregation distortion here. However, at simplex *×* nullplex markers, `polymapR` (and the chi-squared test) assumes a 1:1 segregation ratio, and so cannot accommodate offspring genotypes of 2 and segregation ratios beyond 1:1. This leads them to detect segregation distortion at these SNPs.

Conversely, `polymapR` is more lenient toward “invalid” genotypes, as it only runs its tests on the “valid” genotypes. This leads it to fail to detect segregation distortion at some SNPs where our LRT and Bayes test indicate that there is strong segregation distortion. A few examples of such SNPs are in the first five row in Table S2. At each of these, the tabulated posterior mode genotypes indicate that there are individuals with “invalid” genotypes. E.g., at SNP 12 8929238, genotypes of 3 should be impossible at this simplex *×* nullplex marker, even with double reduction, and so our LRT and Bayes test indicate that there is segregation distortion here. However, `polymapR`’s “valid” genotypes (0 and 1) are at an observed ratio of 106:114, which is close enough to the expected 1:1 ratio that it provides a large *p*-value. The number of “invalid” genotypes is small enough to not be flagged by `polymapR`. Though, we would argue that observing about 11 “invalid” genotypes (for SNP 12 8929238) should flag possible segregation distortion.

## 4 Discussion

We developed new models for the gamete frequencies of tetraploids that incorporate both preferential pairing and double reduction. We used these models to develop likelihood ratio and Bayesian tests for segregation distortion in F1 populations, that optionally account for genotype uncertainty. We demonstrated that our LRT controls Type I error, where competing methods sometimes do not. Our Bayesian test had good performance in simulations, generally supporting the null when the null was true and supporting the alternative when the alternative was true. We demonstrated our methods on a real F1 population of tetraploid blueberries.

Tests for segregation distortion are generally only one part of the quality control pipeline of a study. Indeed, the `polymapR` package’s `checkF1()` function performs various checks, of which segregation distortion is one aspect, and aggregates these results into various quality scores. We imagine that our tests derived here could be similarly used as part of a quality control pipeline, where they can be a drop-in replacement for the standard chi-squared test.

Concerning our new model for gamete frequencies, unfortunately neither the two-parameter nor the three-parameter model are identified when the double reduction rate and the preferential pairing parameter are together. That is, the models are not identified when a parent genotype is *ℓ* = 2. One can see this, for example, by noting that, when *ℓ* = 2, *α* = 1*/*6 and *ξ* = 1*/*3 results in the same gamete frequencies as *α* = 0 and *ξ* = 1*/*9. This means that one cannot estimate *ξ* (or *α* when *ℓ* = 2). However, our model indicates that one need not worry about preferential pairing at loci where *ℓ* = 1 or 3 and can use these loci to estimate the double reduction rate, *α*. We note that though the model is unidentified when *ℓ* = 2, this is not a major issue for our purpose of hypothesis testing. The unidentifiability affects the number of degrees of freedom calculation for the LRTs of Section 2.2, and merely affects the prior distribution over the null parameter space for our Bayesian tests in Section 2.3.

Though the model in Table 2 contains only two parameters, it is not always preferred to that of Table 1 because the range of the parameters in the two-parameter model are dependent. This results from the well-known fact that, under various models, there is an upper bound on the rate of double reduction [Mather, 1935, Huang et al., 2019]. E.g., under the complete equational segregation model, the maximum value of the double reduction rate is 1*/*6 (so 0 ≤ *α* ≤ *β* ≤ 1*/*6). Suppose that the maximum rate is *c*, then we have by Theorem S2 that

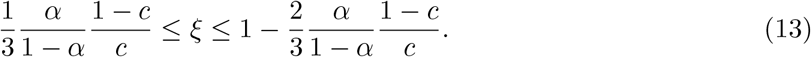

The preferential pairing parameter, *γ*, is interpreted as the frequency of bivalent pairing between chromosomes carrying certain alleles. Since individuals might have different alleles on different subgenomes, this has a few consequences for the broader applicability of our model. First, each parent may contain different alleles on different subgenomes, and so each parent should have their preferential pairing parameter modeled separately (either *γ* or *ξ*). Second, as offspring may have different alleles on different subgenomes, this model will not be persistent across more than one F1 population. Thus, it should not naively be used for simulating multiple generations.

## Acknowledgments

This material is based upon work supported by the National Science Foundation under Grant No. 2132247.

Most analyses were performed using the R statistical language [R Core Team, 2022].

## Data accessibility

The methods described in this paper are implemented in the menbayes package on GitHub: https://github.com/dcgerard/menbayes

All analysis scripts and data needed to reproduce the results of this paper are available on GitHub: https://github.com/dcgerard/mbanalysis

## Supplementary material

Additional figures, tables, and theoretical details are available in the Supplementary Material online.

## Author contributions

DG developed the methodology, wrote the software, implemented the study, and wrote the manuscript.

MT wrote the software, implemented the study, and wrote the manuscript.

LFVF implemented the study and wrote the manuscript.

## Notes

https://github.com/dcgerard/menbayes

https://github.com/dcgerard/mbanalysis

### Competing Interest Statement

The authors have declared no competing interest.

