## Supplementary Material for "Tests for Segregation Distortion in Tetraploid F1 Populations"

**Abstract**

This document contains additional theoretical considerations, derivations, and figures to supplement the manuscript "Tests for Segregation Distortion in Tetraploid F1 Populations".

#### S1 Related work

In this section, we outline prior research on double reduction or preferential pairing in F1 populations. All of these approaches have limitations: (i) none account for both double reduction and preferential pairing at a single biallelic locus simultaneously, (ii) none consider genotype uncertainty, a common issue in polyploid genetics [Gerard et al., 2018, Gerard and Ferrão, 2019], and (iii) most focus on estimating meiotic parameters, rather than testing for segregation distortion, which is our objective.

Many previous approaches have estimated the double reduction rate using gamete frequencies, although the ones mentioned in this paragraph do not consider partial preferential pairing. Fisher and Mather [1943] provides a model for gamete inheritance in the presence of double reduction, but not preferential pairing. This was later generalized to higher ploidies by Huang et al. [2019]. However, neither of these papers provide estimation and testing strategies related to these frequencies. Tai [1982a] and Tai [1982b] estimate the double reduction rate using a complex series of crosses between diploids and tetraploids, but they do not account for preferential pairing, and their scheme was in the context of estimating the quantitative effects of genotypes on phenotypic traits. Haynes and Douches [1993] estimates double reduction, but assumes that one can obtain the gamete genotypes, and also does not account for preferential pairing. Bourke et al. [2015] looked for evidence of double reduction by finding duplex offspring markers in nullplex by simplex crosses but did not provide general methods for more general parental genotypes. Gerard [2022b] estimates double reduction for autopolyploid populations in Hardy-Weinberg equilibrium, but not for F1 populations [see also Gerard, 2022a].

Previous approaches to estimating preferential pairing are not designed for biallelic SNP data or are otherwise limiting. Qu and Hancock [2001] and Bourke et al. [2017] developed methods to estimate the degree of preferential pairing but require simplex by nullplex markers 0 cM apart that are in repulsion linkage. Bourke et al. [2017] can estimate the degree of preferential pairing only when the marker alleles are located on the homologues, but *not* the homoeologues. Cao et al. [2004] also requires simplex by nullplex markers in repulsion linkage but allows for markers to be further than 0 cM apart. The method of Wu et al. [2001] is not designed for biallelic SNP data, assumes equal preferential pairing at both ends of the chromosome, and provides no software implementing the methods. The model of Wu et al. [2001] further assumes that quadrivalent frequencies of less than 2/3 are the result of preferential pairing, an assumption violated in some organisms such as *Solanum tuberosum* [Swaminathan and Howard, 1953] and *Lotus corniculatus* [Fjellstrom et al., 2001]. Olson [1997] only models random chromosome segregation and disomic segregation, not accounting for double reduction and partial preferential pairing. Sun [2020] provides a model and estimation procedure for preferential pairing for any ploidy but does not account for double reduction. All of these methods also assume that genotypes are known without error, an unrealistic assumption in polyploids [Gerard et al., 2018, Gerard and Ferrão, 2019].

Polyploid phasing software often accounts for preferential pairing or double reduction, and so the degree of preferential pairing or the double reduction rate are estimated as a by-product [Zheng et al., 2016, Bourke et al., 2018, Mollinari et al., 2020, Zheng et al., 2021]. However, such methods either do not allow for heterogeneous levels of preferential pairing [Zheng et al., 2016, Bourke et al., 2018, Zheng et al., 2021], or do not accommodate double reduction [Bourke et al., 2018, Mollinari et al., 2020], which we show in Section 4 can be confounded with the effects of preferential pairing.

One paper that does account for both double reduction and preferential pairing in tetraploids is Stift et al. [2008]. This paper provides a general model for chromosome segregation in tetraploids with arbitrary levels of double reduction and preferential pairing. However, the model of Stift et al. [2008] assumes that all chromosomes are uniquely marked, and that gametes can be uniquely genotyped. This is not typically the case for modern biallelic SNP markers. In Section S2, we use the model of Stift et al. [2008] as a starting point to derive a model for gamete frequencies that are agnostic to chromosomal assignment, and may thus be applied to biallelic SNP data.

### S2 A model for gamete frequencies, accounting for double reduction and preferential pairing

Stift et al. [2008] proposed a model for segregation in tetraploids that accounts for both preferential pairing and double reduction. This model assumes that all four chromosomes can be distinguished. However, researchers typically only have biallelic dosage data (i.e. the number of copies of the alternative allele for an individual at a locus) [Gerard et al., 2018, Gerard and Ferrão, 2019] where each chromosome cannot be separately determined. Here, we will modify the model of Stift et al. [2008] over chromosome assignment to derive a model for segregation at biallelic loci in tetraploids.

We begin by describing the model of Stift et al. [2008]. Let $c_1$, $c_2$, $c_3$, and $c_4$ denote the four chromosomes of a tetraploid individual. The model of Stift et al. [2008] has three parameters. Let $\tau$ be the proportion of quadrivalent pairing, let $\beta$ be the probability of double reduction given quadrivalent pairing, so $\alpha = \tau\beta$ is the double reduction rate, and let $(\delta_1, \delta_2, \delta_3)$ be the rates of different bivalent pairings, where $\delta_1$ is the probability of $c_1$ and $c_2$ pairing together given bivalent formation, $\delta_2$ is the probability of $c_1$ and $c_3$ pairing together given bivalent formation, and $\delta_3$ is the probability of $c_1$ and $c_4$ pairing together given bivalent formation.. Note that $\delta_1 + \delta_2 + \delta_3 = 1$ and, for identifiability reasons, Stift et al. [2008] constrain $\delta_1\delta_2\delta_3 = 0$ (at least one pairing has zero probability), but we don't make this identifying assumption. Then the model of Stift et al. [2008]

states the probability of each gamete to be

$$\begin{pmatrix} \Pr(c_1c_1) \\ \Pr(c_2c_2) \\ \Pr(c_3c_3) \\ \Pr(c_4c_4) \\ \Pr(c_1c_2) \\ \Pr(c_1c_3) \\ \Pr(c_1c_4) \\ \Pr(c_3c_4) \\ \Pr(c_2c_4) \\ \Pr(c_2c_3) \end{pmatrix} = \begin{pmatrix} 0 \\ 0 \\ 0 \\ 0 \\ 1/6 \\ 1/6 \\ 1/6 \\ 1/6 \\ 1/6 \\ 1/6 \end{pmatrix} \tau + \begin{pmatrix} 1/4 \\ 1/4 \\ 1/4 \\ 1/4 \\ -1/6 \\ -1/6 \\ -1/6 \\ -1/6 \\ -1/6 \\ -1/6 \end{pmatrix} \beta\tau + (1-\tau) \begin{pmatrix} 0 & 0 & 0 \\ 0 & 0 & 0 \\ 0 & 0 & 0 \\ 0 & 0 & 0 \\ 0 & 1/4 & 1/4 \\ 1/4 & 0 & 1/4 \\ 1/4 & 1/4 & 0 \\ 0 & 1/4 & 1/4 \\ 1/4 & 0 & 1/4 \\ 1/4 & 1/4 & 0 \end{pmatrix} \begin{pmatrix} \delta_1 \\ \delta_2 \\ \delta_3 \end{pmatrix}. \tag{S1}$$

More concisely, equation (S1) may be written as

$$Pr(c_1c_1) = Pr(c_2c_2) = Pr(c_3c_3) = Pr(c_4c_4) = \frac{1}{4}\beta\tau, \tag{S2}$$

$$Pr(c_1c_2) = Pr(c_3c_4) = \frac{1}{6}\tau(1-\beta) + \frac{1}{4}(1-\tau)(\delta_2+\delta_3), \tag{S3}$$

$$Pr(c_1c_3) = Pr(c_2c_4) = \frac{1}{6}\tau(1-\beta) + \frac{1}{4}(1-\tau)(\delta_1+\delta_3), \tag{S4}$$

$$Pr(c_1c_4) = Pr(c_2c_3) = \frac{1}{6}\tau(1-\beta) + \frac{1}{4}(1-\tau)(\delta_1+\delta_2). \tag{S5}$$

We will now modify model (S2)–(S5) to biallelic dosage data. Let $\ell \in \{0,1,2,3,4\}$ be the dosage of the parent at a locus, and let $x \in \{0,1,2\}$ be the random variable of the number of alternative alleles that the parent sends to an offspring. We assume that we do not know which chromosomes contain the alternative and reference alleles. Then, for a parental dosage of $\ell = 0$, none of the chromosomes have the reference allele, and we have

$$\begin{aligned} Pr(x=0|\ell=0) &= Pr(c_1c_2) + Pr(c_1c_3) + Pr(c_1c_4) + Pr(c_2c_3) + Pr(c_2c_4) \\ &\quad + Pr(c_3c_4) + Pr(c_1c_1) + Pr(c_2c_2) + Pr(c_3c_3) + Pr(c_4c_4) \\ &= 1 \end{aligned} \tag{S6}$$

$$Pr(x=1|\ell=0) = Pr(x=2|\ell=0) = 0. \tag{S7}$$

By symmetry, we have

$$Pr(x=0|\ell=4) = Pr(x=1|\ell=4) = 0 \tag{S8}$$

$$Pr(x=2|\ell=4) = 1. \tag{S9}$$

For a parental dosage of $\ell = 1$, for the moment allow $c_1$ to carry the $\mathsf{A}$ allele, and $c_2$, $c_3$, and $c_4$ to carry the $\mathsf{a}$ allele. Then

$$Pr(x=0|\ell=1) = Pr(c_2c_3) + Pr(c_2c_4) + Pr(c_3c_4) + Pr(c_2c_2) + Pr(c_3c_3) + Pr(c_4c_4) \tag{S10}$$

$$= \frac{1}{2}\tau(1-\beta) + \frac{1}{4}(1-\tau)(\delta_1+\delta_2+\delta_1+\delta_3+\delta_2+\delta_3) + \frac{3}{4}\beta\tau \tag{S11}$$

$$= \frac{1}{2}\tau(1-\beta) + \frac{1}{2}(1-\tau) + \frac{3}{4}\beta\tau \tag{S12}$$

$$= \frac{1}{2} + \frac{1}{4}\beta\tau. \tag{S13}$$

$$Pr(x=1|\ell=1) = Pr(c_1c_2) + Pr(c_1c_3) + Pr(c_1c_4) \tag{S14}$$

$$= \frac{1}{2}\tau(1-\beta) + \frac{1}{4}(1-\tau)(\delta_2+\delta_3+\delta_1+\delta_3+\delta_1+\delta_2) \tag{S15}$$

$$= \frac{1}{2}\tau(1-\beta) + \frac{1}{2}(1-\tau) \tag{S16}$$

$$= \frac{1}{2} - \frac{1}{2}\beta\tau \tag{S17}$$

$$Pr(x=2|\ell=1) = Pr(c_1c_1) = \frac{1}{4}\beta\tau. \tag{S18}$$

This probability is independent of the labeling for which chromosome carries the A allele, and so is the probability distribution of a gamete dosage given the parental dosage. By symmetry, we have

$$Pr(x=0|\ell=3) = \frac{1}{4}\beta\tau, \tag{S19}$$

$$Pr(x=1|\ell=3) = \frac{1}{2} - \frac{1}{2}\beta\tau \tag{S20}$$

$$Pr(x=2|\ell=3) = \frac{1}{2} + \frac{1}{4}\beta\tau. \tag{S21}$$

For a parental dosage of $\ell = 2$, for the moment allow $c_1$ and $c_2$ to carry the A allele, and $c_3$ and $c_4$ to carry the a allele. Then

$$Pr(x=0|\ell=2) = Pr(c_3c_4) + Pr(c_3c_3) + Pr(c_4c_4) \tag{S22}$$

$$= \frac{1}{6}\tau(1-\beta) + \frac{1}{4}(1-\tau)(\delta_2+\delta_3) + \frac{1}{2}\beta\tau \tag{S23}$$

$$= \frac{1}{3}\beta\tau + \frac{1}{6}\tau + \frac{1}{4}(1-\tau)(1-\delta_1). \tag{S24}$$

By symmetry we have

$$Pr(x=2|\ell=2) = \frac{1}{3}\beta\tau + \frac{1}{6}\tau + \frac{1}{4}(1-\tau)(1-\delta_1). \tag{S25}$$

Finally,

$$Pr(x=1|\ell=2) = Pr(c_1c_3) + Pr(c_1c_4) + Pr(c_2c_3) + Pr(c_2c_4) \tag{S26}$$

$$= \frac{2}{3}\tau(1-\beta) + \frac{1}{4}(1-\tau)(\delta_1+\delta_3+\delta_1+\delta_2+\delta_1+\delta_2+\delta_1+\delta_3) \tag{S27}$$

$$= -\frac{2}{3}\beta\tau + \frac{2}{3}\tau + \frac{1}{2}(1-\tau)(1+\delta_1). \tag{S28}$$

Notice that $\delta_1$ is the probability that the chromosomes that share the same alleles will pair ($c_1$ with $c_2$ and $c_3$ with $c_4$). To make these probabilities independent of the labeling of the chromosomes, we set $\gamma$ to be the probability that the chromosomes that share the same alleles will pair, obtaining

$$Pr(x=0|\ell=2) = Pr(x=2|\ell=2) = \frac{1}{3}\beta\tau + \frac{1}{6}\tau + \frac{1}{4}(1-\tau)(1-\gamma), \text{ and} \tag{S29}$$

$$Pr(x = 1|\ell = 2) = -\frac{2}{3}\beta\tau + \frac{2}{3}\tau + \frac{1}{2}(1 - \tau)(1 + \gamma), \tag{S30}$$

where $\gamma \in \{\delta_1, \delta_2, \delta_3\}$. We summarize the probabilities of $Pr(x|\ell)$ in Table 1.

The model in 1 contains three parameters. However, we can reduce it down to two parameters. Let $\eta$ be the probability of quadrivalent formation given no double reduction. That is, using Bayes rule,

$$\eta = \frac{(1 - \beta)\tau}{(1 - \beta)\tau + (1 - \tau)}. \tag{S31}$$

Let $\xi$ be a convex combination between $\gamma$ and 1/3, weighted by $\eta$

$$\xi = \eta\frac{1}{3} + (1 - \eta)\gamma. \tag{S32}$$

This $\xi$ parameter measures the degree of preferential pairing, where deviations from 1/3 represent deviations from autopolyploidy, while deviations from 1 or 0 represent deviations from allopolyploidy. Finally, let $\alpha$ be the marginal probability of double reduction

$$\alpha = \beta\tau. \tag{S33}$$

Then, using the parameters $\alpha$ and $\xi$, we may re-write the probability distributions in Table 1 as Table 2 (Theorem S1).

**Theorem S1.** *Let $\xi$ and $\alpha$ be as defined in (S32) and (S33), respectively. Then the probability distributions in Tables 1 and 2 are equivalent.*

*Proof.* The correspondence between $Pr(x|\ell = 0)$, $Pr(x|\ell = 1)$, $Pr(x|\ell = 3)$, and $Pr(x|\ell = 4)$ between the two tables is obvious. It suffices to show that

$$\frac{1}{2}(1 - \alpha)(1 + \xi) = -\frac{2}{3}\beta\tau + \frac{2}{3}\tau + \frac{1}{2}(1 - \tau)(1 + \gamma), \tag{S34}$$

as the other equalities from $Pr(x|\ell = 2)$ would follow by the sum-to-one constraint. From the left-hand side of (S34), we have

$$\frac{1}{2}(1 - \alpha)(1 + \xi) = \frac{1}{2}(1 - \beta\tau)(1 + \eta\frac{1}{3} + (1 - \eta)\gamma) \tag{S35}$$

$$= \frac{1}{2}(1 - \beta\tau)\left(1 + \frac{\frac{1}{3}(1 - \beta)\tau}{(1 - \beta)\tau + (1 - \tau)} + \frac{(1 - \tau)\gamma}{(1 - \beta)\tau + (1 - \tau)}\right) \tag{S36}$$

$$= \frac{1}{2}(1 - \beta\tau)\left(\frac{(1 - \beta)\tau + (1 - \tau) + \frac{1}{3}(1 - \beta)\tau + (1 - \tau)\gamma}{(1 - \beta)\tau + (1 - \tau)}\right) \tag{S37}$$

$$= \frac{1}{2}(1 - \beta\tau)\left(\frac{\frac{4}{3}(1 - \beta)\tau + (1 - \tau)(1 + \gamma)}{(1 - \beta)\tau + (1 - \tau)}\right) \tag{S38}$$

$$= \frac{1}{2}(1 - \beta\tau)\left(\frac{\frac{4}{3}(1 - \beta)\tau + (1 - \tau)(1 + \gamma)}{1 - \beta\tau}\right) \tag{S39}$$

$$= \frac{2}{3}(1-\beta)\tau + \frac{1}{2}(1-\tau)(1+\gamma) \tag{S40}$$

$$= -\frac{2}{3}\beta\tau + \frac{2}{3}\tau + \frac{1}{2}(1-\tau)(1+\gamma). \tag{S41}$$

□

Though we have managed to reduce the number of parameters from three to two, the resulting parameterization induces a dependence on the range of possible value of $\xi$ given a value of $\alpha$ (Theorem S2).

**Theorem S2.** *Suppose that $0 \leq \beta \leq c$. Then, for a given value of $\alpha$, we have*

$$\frac{1}{3}\frac{\alpha}{1-\alpha}\frac{1-c}{c} \leq \xi \leq 1 - \frac{2}{3}\frac{\alpha}{1-\alpha}\frac{1-c}{c}. \tag{S42}$$

*Proof.* We may rewrite $\xi$ as

$$\xi = \frac{1}{3}\frac{\tau-\beta\tau}{1-\beta\tau} + \frac{1-\tau}{1-\beta\tau}\gamma \tag{S43}$$

$$= \frac{1}{3}\frac{\tau-\alpha}{1-\alpha} + \frac{1-\tau}{1-\alpha}\gamma. \tag{S44}$$

Since $\tau = \alpha/\beta$ (and $\alpha \leq \beta$), we have, for a given value of $\alpha$, that $\alpha/c \leq \tau \leq 1$.

To find the minimum value of $\xi$, we minimize (S44) over $\tau$ and $\gamma$. This minimum occurs at $\gamma = 0$ and $\tau = \alpha/c$, yielding the lower bound of (S42). To find the maximum value of $\xi$, we maximize (S44) over $\tau$ and $\gamma$. This maximum occurs at $\gamma = 1$ and $\tau = \alpha/c$, yielding the upper bound of (S42). □

#### S3 Generalization of Fisher and Mather [1943]

Here, we show that our models for the gamete frequencies in Table 2 (and, thus, Table 1) are generalizations of those in Table 9 from Fisher and Mather [1943]. The model from Fisher and Mather [1943] incorporates double reduction, but not preferential pairing, and we show that setting $\xi = 1/3$ results in the same gamete frequencies as those in Table 9 from Fisher and Mather [1943]. It is trivial to check the equivalence between Table 2 and Table 9 from Fisher and Mather [1943] for the parental genotypes $\ell = 0, 1, 3, 4$, so we just consider $\ell = 2$. For our tetraploid model, we have that $\xi$ is the probability of pairing configuration A-A:a-a, and $1-\xi$ is the probability of pairing configuration A-a:A-a. Labeling $c_1, c_2, c_3, c_4$ as the four chromosomes, and allowing $c_1$ and $c_2$ to carry the A alleles, the possible pairings are $c_1$-$c_2$:$c_3$-$c_4$ = A-A;a-a, $c_1$-$c_3$:$c_2$-$c_4$ = A-a:A-a, and $c_1$-$c_4$:$c_2$-$c_3$ = A-a:A-a. Thus, under polysomic inheritance, A-A:a-a occurs with probability $\xi = 1/3$. We can plug this in to get the gamete frequencies under polysomic inheritance

$$\Pr(x = 0|\ell = 2, \xi = 1/3) = \frac{1}{2}\alpha + \frac{1}{4}(1-\frac{1}{3})(1-\alpha) \tag{S45}$$

$$= \frac{1}{6} + \frac{1}{3}\alpha, \tag{S46}$$

which one can check is the same as the value as Table 9 from Fisher and Mather [1943]. The other values for $\ell = 2$ follow from symmetry and the sum-to-one constraint.

#### S4 Supplementary tables, figures, and procedures

| Condition | Method | .05 | .01 | .001 | .0001 | .00001 |
|---|---|---|---|---|---|---|
| n=20,rd=10 | Chisq | 0.98 | 0.96 | 0.90 | 0.84 | 0.80 |
| n=20,rd=10 | LRT | 0.55 | 0.34 | 0.15 | 0.04 | 0.01 |
| n=20,rd=10 | polymapR | 0.80 | 0.66 | 0.43 | 0.27 | 0.27 |
| n=20,rd=Inf | Chisq | 0.99 | 0.96 | 0.91 | 0.87 | 0.79 |
| n=20,rd=Inf | LRT | 0.67 | 0.46 | 0.22 | 0.11 | 0.04 |
| n=20,rd=Inf | polymapR | 0.78 | 0.64 | 0.47 | 0.28 | 0.28 |
| n=200,rd=10 | Chisq | 1.00 | 1.00 | 1.00 | 1.00 | 1.00 |
| n=200,rd=10 | LRT | 1.00 | 1.00 | 1.00 | 1.00 | 1.00 |
| n=200,rd=10 | polymapR | 1.00 | 1.00 | 1.00 | 1.00 | 1.00 |
| n=200,rd=Inf | Chisq | 1.00 | 1.00 | 1.00 | 1.00 | 1.00 |
| n=200,rd=Inf | LRT | 1.00 | 1.00 | 1.00 | 1.00 | 1.00 |
| n=200,rd=Inf | polymapR | 1.00 | 1.00 | 1.00 | 1.00 | 1.00 |

Table S1: Power for different methods (Method) at different samples sizes and read-depth (Condition) at different significance levels (.05 through .00001). The methods considered are the standard chi-squared test (Chisq), the likelihood ratio test of Section 2.2 (LRT), and the polymapR test of Section 1.1.

| SNP | $\ell_1$ | $\ell_2$ | Bayes | LRT | polymapR | Observed | Expected |
|---|---|---|---|---|---|---|---|
| 12_8929238 | 0 | 1 | -35 | 3.9e-22 | 0.97 | (105,113,11,11,0) | (130,100,10,0,0) |
| 11_32341161 | 2 | 4 | -16 | 1.8e-15 | 0.81 | (0,8,35,163,34) | (0,0,38.8,162.3,38.8) |
| 6_15037920 | 0 | 2 | -16 | 6.2e-14 | 0.96 | (36,153,44,7,0) | (42.7,154.6,42.7,0,0) |
| 6_14723914 | 4 | 2 | -13 | 4.8e-12 | 0.84 | (0,7,40,156,37) | (0,0,42.1,155.8,42.1) |
| 17_9734135 | 0 | 3 | -6.5 | 2.1e-10 | 0.83 | (23,102,108,6,1) | (10,100,130,0,0) |
| 1_1019768 | 1 | 0 | 11 | 0.97 | 0.0016 | (132,100,8,0,0) | (130,100,10,0,0) |
| 22_23904806 | 0 | 1 | 11 | 0.88 | 0.002 | (134,97,9,0,0) | (130,100,10,0,0) |
| 2_12659258 | 1 | 0 | 11 | 0.99 | 0.002 | (131,101,8,0,0) | (130,100,10,0,0) |
| 2_12659201 | 1 | 0 | 11 | 0.98 | 0.002 | (131,101,8,0,0) | (130,100,10,0,0) |
| 12_30140888 | 0 | 1 | 9.9 | 0.89 | 0.0031 | (132,97,11,0,0) | (130,100,10,0,0) |

Table S2: In the first five SNPs (plotted in Figure S11), the LRT indicates segregation distortion while the polymapR test indicates no segregation distortion, while in the last five SNPs (plotted in Figure S12) the LRT indicates no segregation distortion while the polymapR test indicates segregation distortion. Parent genotypes ($\ell_1$ and $\ell_2$) are listed, along with the log Bayes factors ("Bayes"), the LRT $p$-values ("LRT"), the polyampR $p$-values ("polymapR"), the observed counts when tabulating posterior mode genotypes ("Observed"), and the expected counts based on the maximum likelihood estimates of $\alpha$ and the $\xi$'s ("Expected").

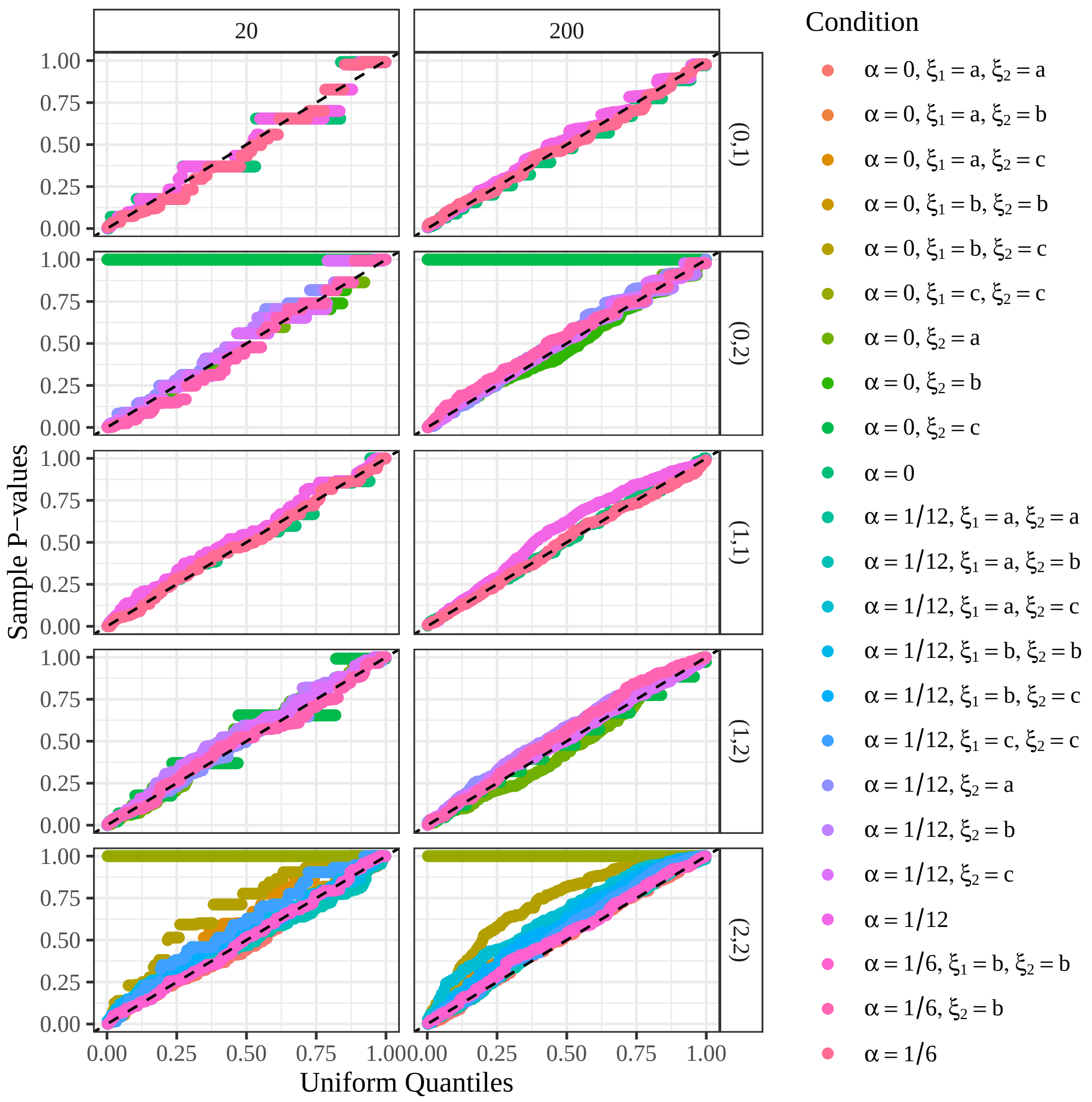


Figure S1: Quantile-quantile plots against the uniform distribution of the $p$-values of the likelihood ratio test of Section 2.2, when using known genotypes, from the null simulations in Section 3.1. Since the null is true, the points should lie either near or above the $y = x$ line (black dashed line). Points that are above the $y = x$ line are conservative, points that are below the $y = x$ line are anti-conservative. Row-facets index parent genotypes $(\ell_1, \ell_2)$, and column-facets index sample size. Color indexes different values of the double reduction rate, $\alpha$, and the preferential pairing parameters of the two parents, $\xi_1$ and $\xi_2$. A preferential pairing value of "a" indicates the lower bound in (S42), a value of "b" indicates 1/3, and a value of "c" indicates the upper bound in (S42). Preferential pairing only affects offspring genotype frequencies when the parent genotype is 2, and so an omission of $\xi_1$ or $\xi_2$ from the color legend indicates a scenario where the corresponding parent genotype is not 2.

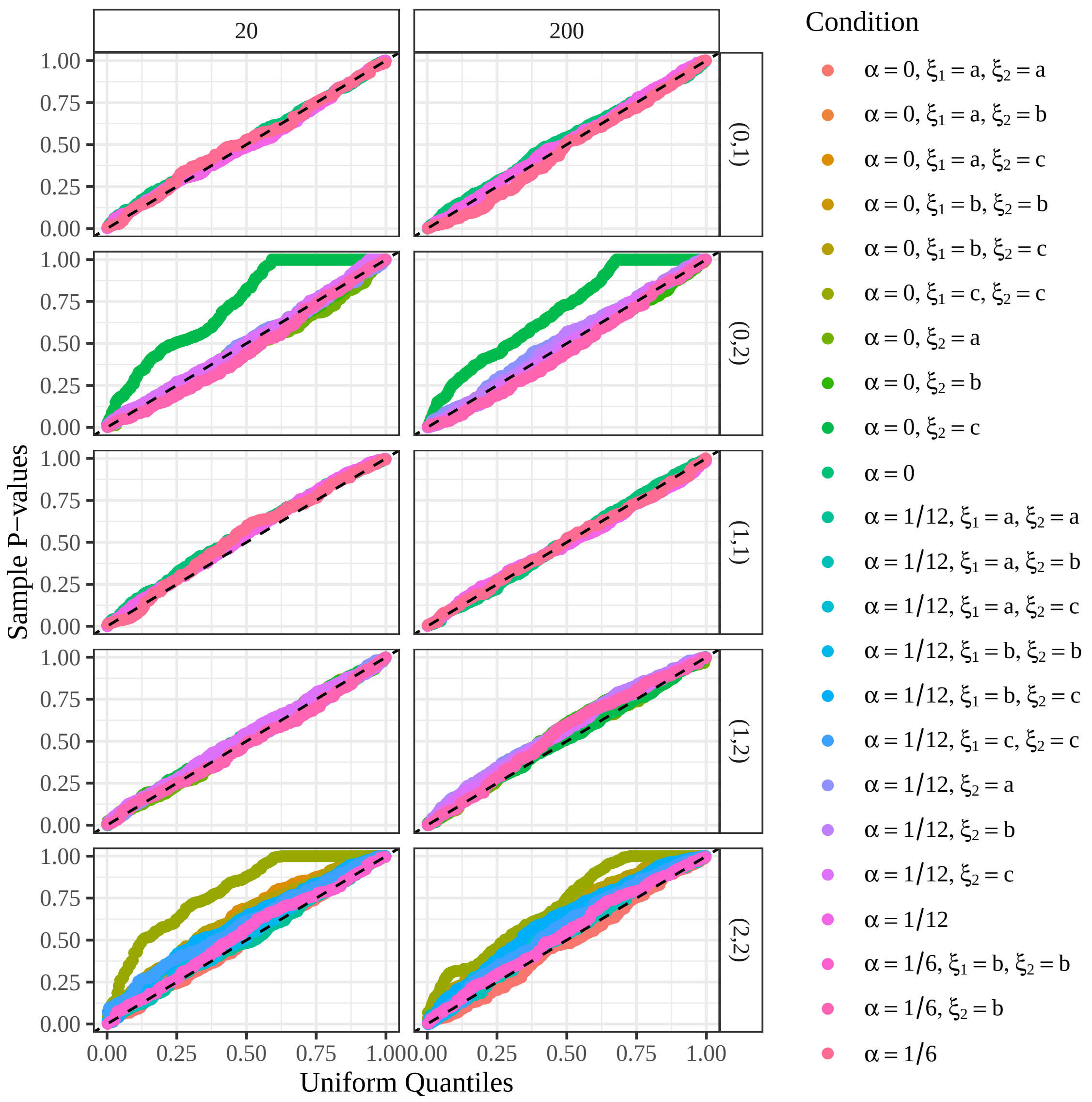


Figure S2: Quantile-quantile plots against the uniform distribution of the $p$-values of the likelihood ratio test of Section 2.2, using genotype likelihoods derived from read-counts simulated at a read-depth of 10, from the null simulations in Section 3.1. Since the null is true, the points should lie either near or above the $y = x$ line (black dashed line). Points that are above the $y = x$ line are conservative, points that are below the $y = x$ line are anti-conservative. Row-facets index parent genotypes $(\ell_1, \ell_2)$, and column-facets index sample size. Color indexes different values of the double reduction rate, $\alpha$, and the preferential pairing parameters of the two parents, $\xi_1$ and $\xi_2$. A preferential pairing value of "a" indicates the lower bound in (S42), a value of "b" indicates 1/3, and a value of "c" indicates the upper bound in (S42). Preferential pairing only affects offspring genotype frequencies when the parent genotype is 2, and so an omission of $\xi_1$ or $\xi_2$ from the color legend indicates a scenario where the corresponding parent genotype is not 2.

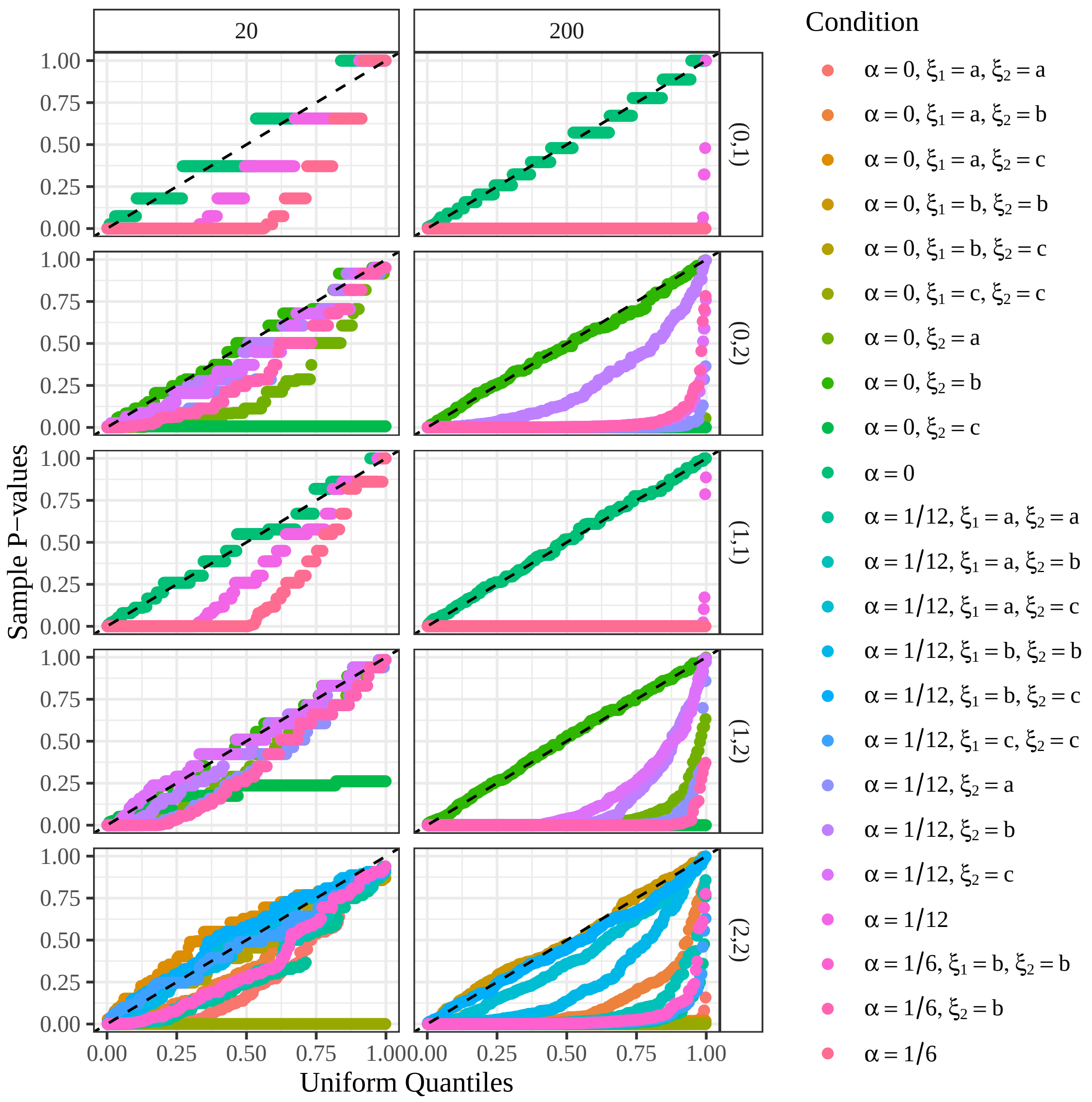


Figure S3: Quantile-quantile plots against the uniform distribution of the $p$-values of the standard chi-squared test for segregation distortion, when using known genotypes, from the null simulations in Section 3.1. Since the null is true, the points should lie either near or above the $y = x$ line (black dashed line). Points that are above the $y = x$ line are conservative, points that are below the $y = x$ line are anti-conservative. Row-facets index parent genotypes $(\ell_1, \ell_2)$, and column-facets index sample size. Color indexes different values of the double reduction rate, $\alpha$, and the preferential pairing parameters of the two parents, $\xi_1$ and $\xi_2$. A preferential pairing value of "a" indicates the lower bound in (S42), a value of "b" indicates 1/3, and a value of "c" indicates the upper bound in (S42). Preferential pairing only affects offspring genotype frequencies when the parent genotype is 2, and so an omission of $\xi_1$ or $\xi_2$ from the color legend indicates a scenario where the corresponding parent genotype is not 2.

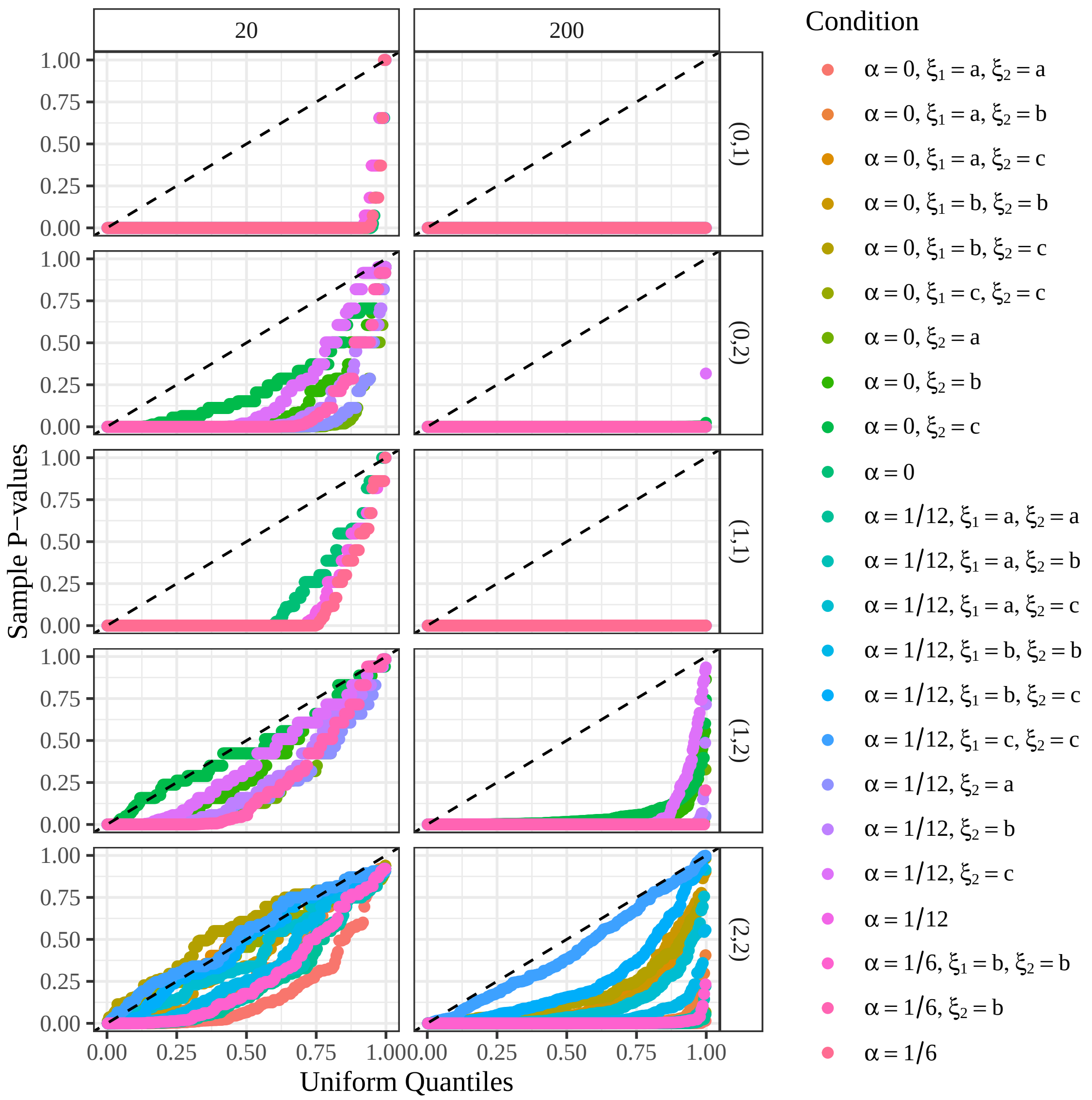


Figure S4: Quantile-quantile plots against the uniform distribution of the $p$-values of the standard chi-squared test for segregation distortion, using genotype likelihoods derived from read-counts simulated at a read-depth of 10, and tabulating posterior mode genotypes, from the null simulations in Section 3.1. Since the null is true, the points should lie either near or above the $y = x$ line (black dashed line). Points that are above the $y = x$ line are conservative, points that are below the $y = x$ line are anti-conservative. Row-facets index parent genotypes $(\ell_1, \ell_2)$, and column-facets index sample size. Color indexes different values of the double reduction rate, $\alpha$, and the preferential pairing parameters of the two parents, $\xi_1$ and $\xi_2$. A preferential pairing value of "a" indicates the lower bound in (S42), a value of "b" indicates 1/3, and a value of "c" indicates the upper bound in (S42). Preferential pairing only affects offspring genotype frequencies when the parent genotype is 2, and so an omission of $\xi_1$ or $\xi_2$ from the color legend indicates a scenario where the corresponding parent genotype is not 2.

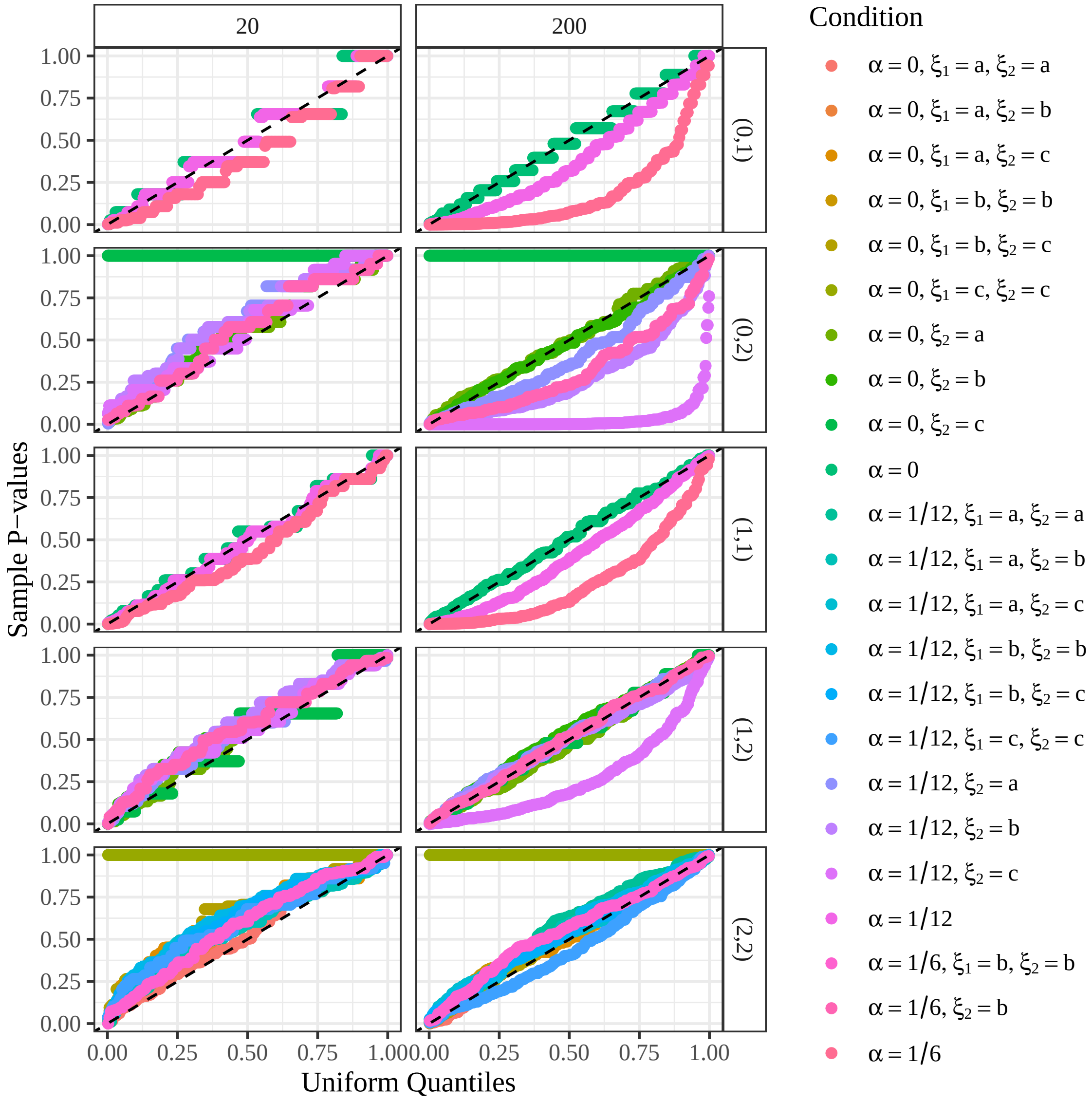


Figure S5: Quantile-quantile plots against the uniform distribution of the $p$-values of the polymapR approach of Section 1.1 [Bourke et al., 2018], when using known genotypes, from the null simulations in Section 3.1. Since the null is true, the points should lie either near or above the $y = x$ line (black dashed line). Points that are above the $y = x$ line are conservative, points that are below the $y = x$ line are anti-conservative. Row-facets index parent genotypes $(\ell_1, \ell_2)$, and column-facets index sample size. Color indexes different values of the double reduction rate, $\alpha$, and the preferential pairing parameters of the two parents, $\xi_1$ and $\xi_2$. A preferential pairing value of "a" indicates the lower bound in (S42), a value of "b" indicates 1/3, and a value of "c" indicates the upper bound in (S42). Preferential pairing only affects offspring genotype frequencies when the parent genotype is 2, and so an omission of $\xi_1$ or $\xi_2$ from the color legend indicates a scenario where the corresponding parent genotype is not 2.

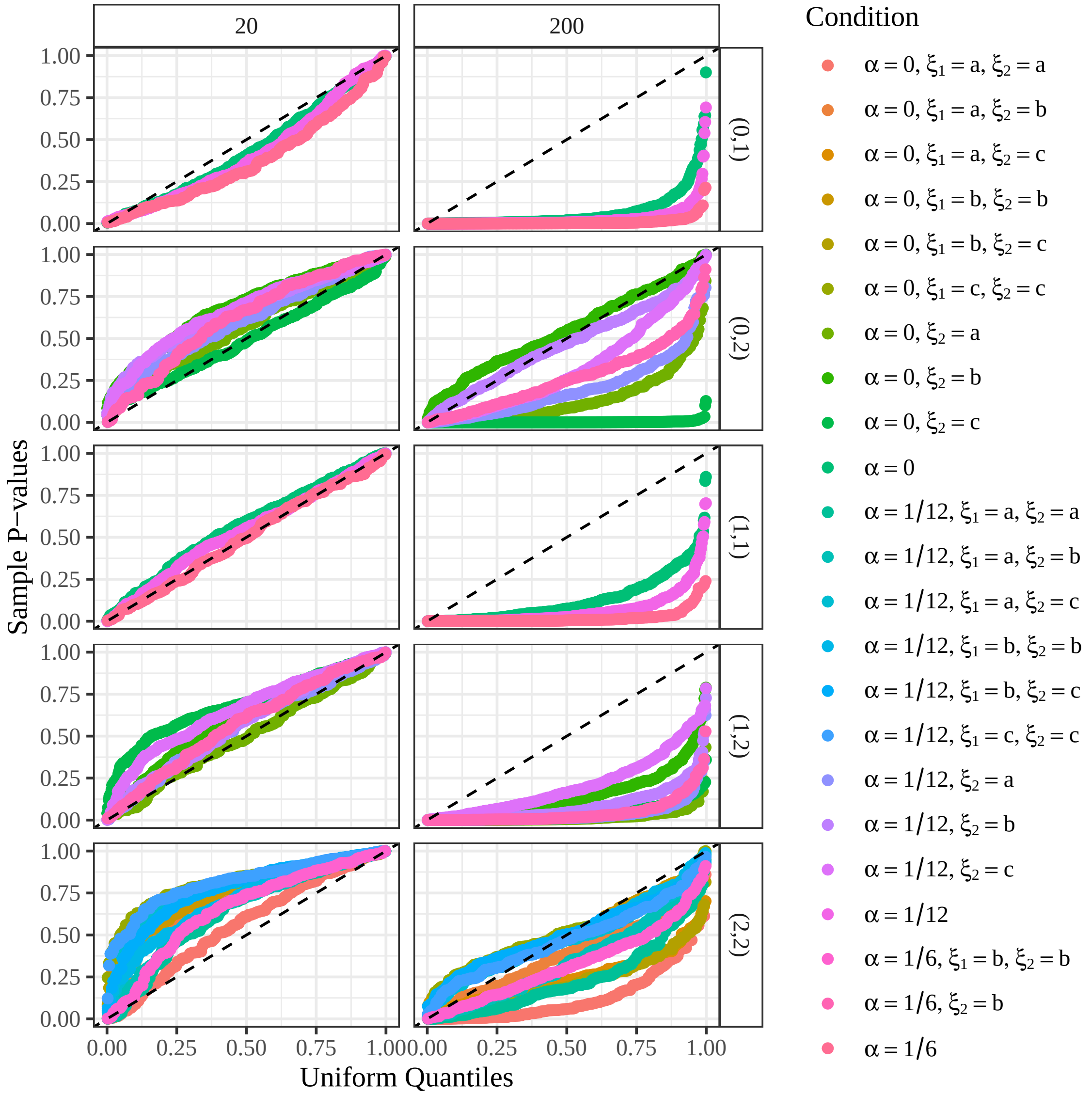


Figure S6: Quantile-quantile plots against the uniform distribution of the $p$-values of the polymapR approach of Section 1.1 [Bourke et al., 2018], using genotype likelihoods derived from read-counts simulated at a read-depth of 10, from the null simulations in Section 3.1. Since the null is true, the points should lie either near or above the $y = x$ line (black dashed line). Points that are above the $y = x$ line are conservative, points that are below the $y = x$ line are anti-conservative. Row-facets index parent genotypes $(\ell_1, \ell_2)$, and column-facets index sample size. Color indexes different values of the double reduction rate, $\alpha$, and the preferential pairing parameters of the two parents, $\xi_1$ and $\xi_2$. A preferential pairing value of "a" indicates the lower bound in (S42), a value of "b" indicates 1/3, and a value of "c" indicates the upper bound in (S42). Preferential pairing only affects offspring genotype frequencies when the parent genotype is 2, and so an omission of $\xi_1$ or $\xi_2$ from the color legend indicates a scenario where the corresponding parent genotype is not 2.

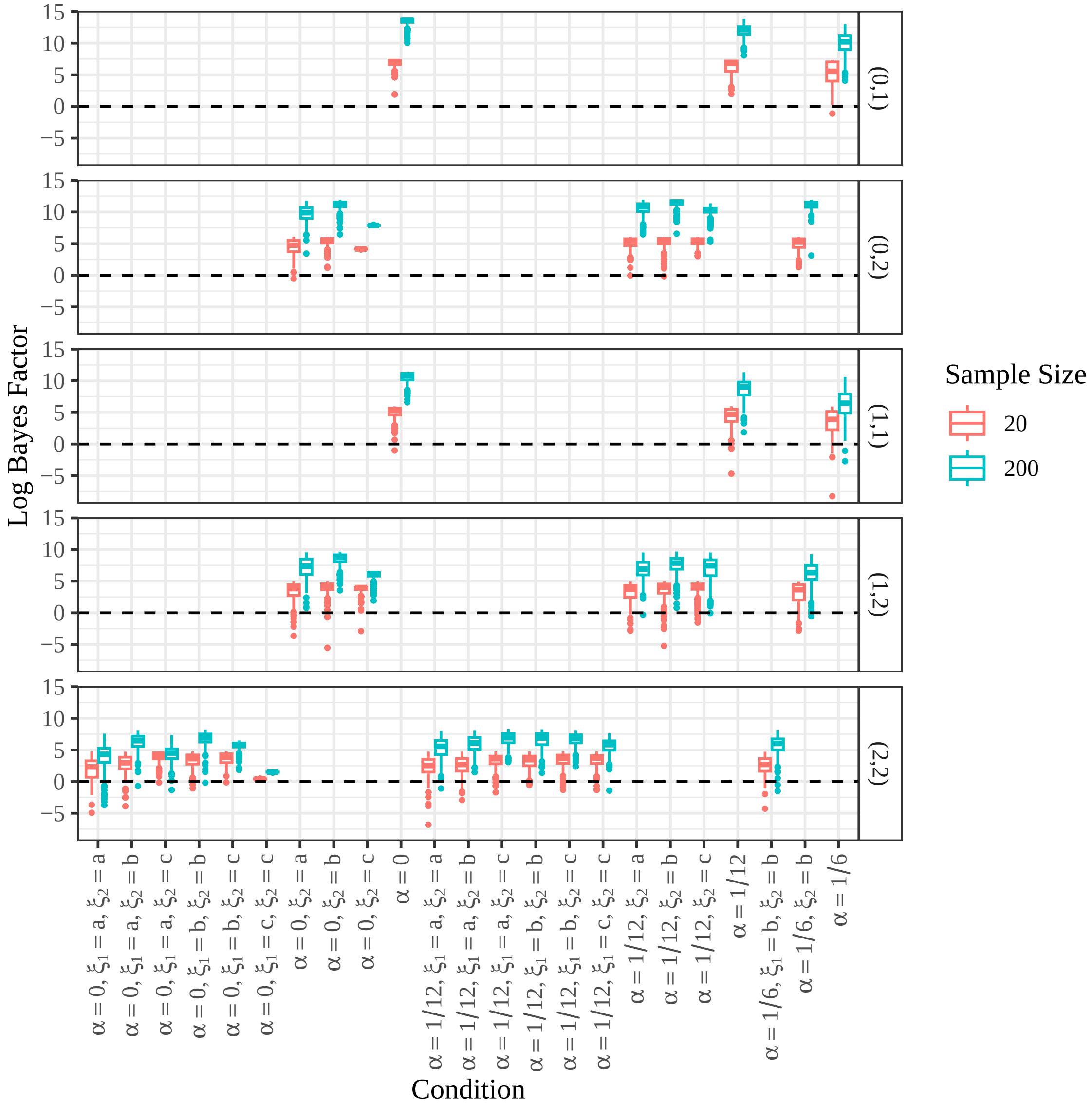


Figure S7: Box plots of the log Bayes factors from the Bayes tests of Section 2.3, when using known genotypes, from the null simulations in Section 3.1. Since the null is true, the log Bayes factors should be mostly above 0 (horizontal black dashed line). Row-facets index parent genotypes $(\ell_1, \ell_2)$, and color indexes sample size. The $x$-axis indexes different values of the double reduction rate, $\alpha$, and the preferential pairing parameters of the two parents, $\xi_1$ and $\xi_2$. A preferential pairing value of "a" indicates the lower bound in (S42), a value of "b" indicates 1/3, and a value of "c" indicates the upper bound in (S42). Preferential pairing only affects offspring genotype frequencies when the parent genotype is 2, and so an omission of $\xi_1$ or $\xi_2$ from the $x$-axis labels indicates a scenario where the corresponding parent genotype is not 2.

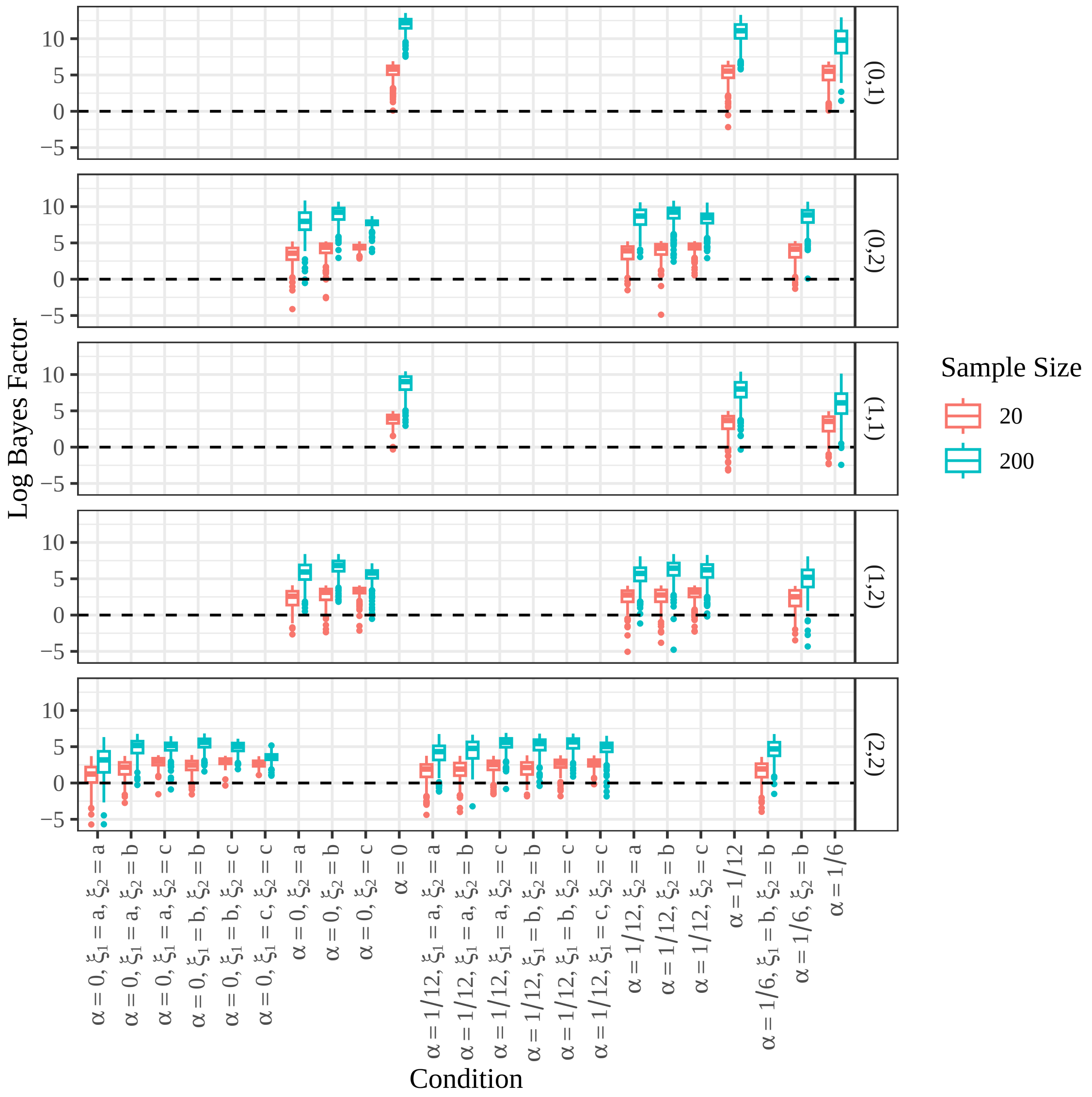


Figure S8: Box plots of the log Bayes factors from the Bayes tests of Section 2.3, using genotype likelihoods derived from read-counts simulated at a read-depth of 10, from the null simulations in Section 3.1. Since the null is true, the log Bayes factors should be mostly above 0 (horizontal black dashed line). Row-facets index parent genotypes $(\ell_1, \ell_2)$, and color indexes sample size. The $x$-axis indexes different values of the double reduction rate, $\alpha$, and the preferential pairing parameters of the two parents, $\xi_1$ and $\xi_2$. A preferential pairing value of “a” indicates the lower bound in (S42), a value of “b” indicates 1/3, and a value of “c” indicates the upper bound in (S42). Preferential pairing only affects offspring genotype frequencies when the parent genotype is 2, and so an omission of $\xi_1$ or $\xi_2$ from the $x$-axis labels indicates a scenario where the corresponding parent genotype is not 2.

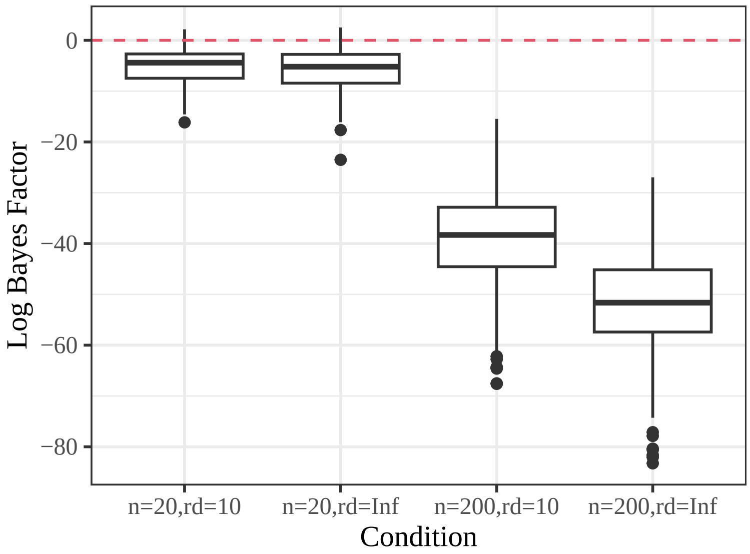


Figure S9: Box plots of the log Bayes factors ($y$-axis) from the Bayesian method of Section 2.3 at different sample sizes ($n$) and different read-depths (rd). Negative values indicate support for the alternative, and these simulations were run when the alternative was true (Section 3.2), and so more negative values indicate superior performance.

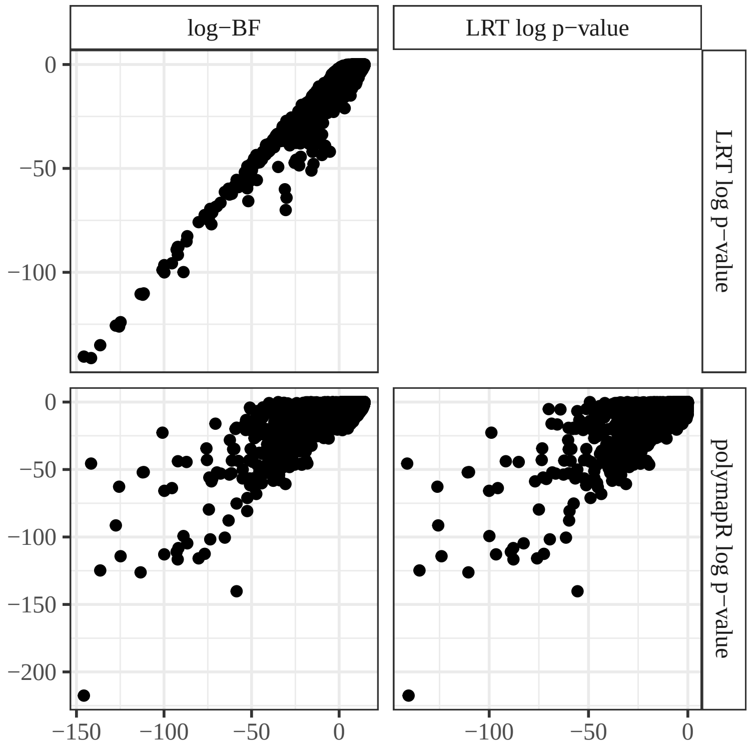


Figure S10: Pairs plot of the log Bayes factors from the Bayes test from Section 2.3, and the log $p$-values from the LRT of Section 2.2 and the polymapR test of Section 1.1. The Bayes test and LRT provide more concordant results.

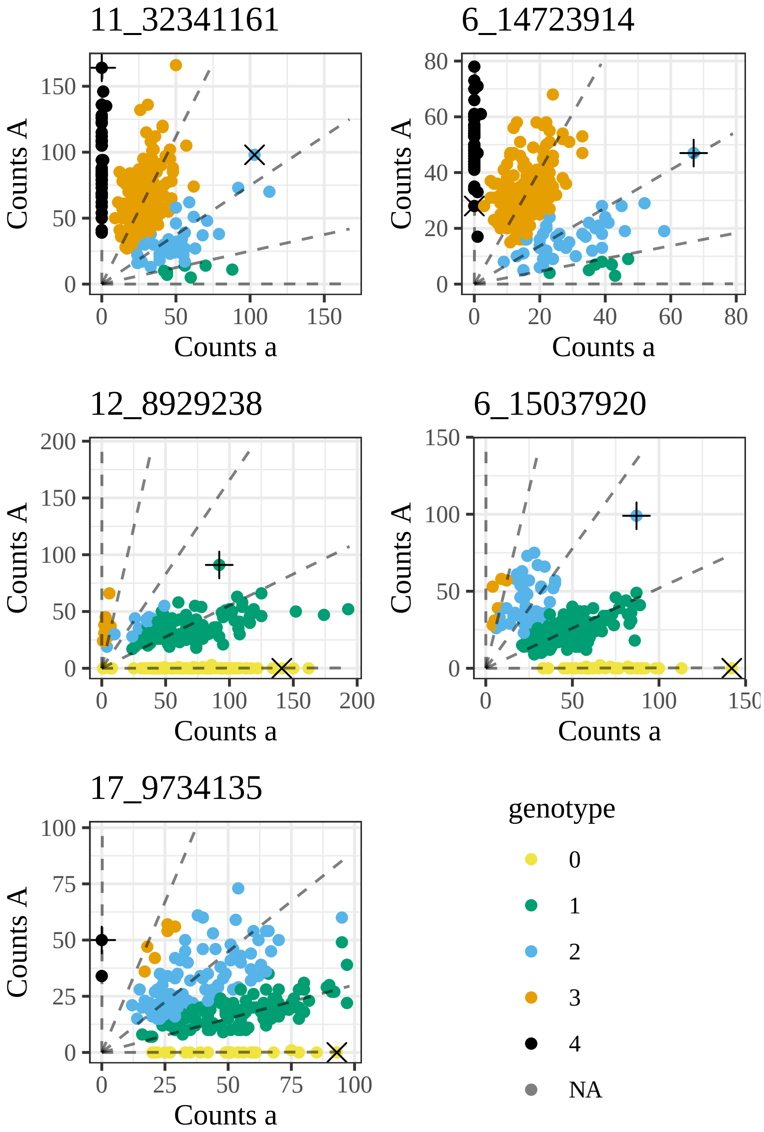


Figure S11: Genotype plots [Gerard et al., 2018] of five SNPs (titles) where the LRT indicates strong segregation distortion, but polymapR indicates no segregation distortion. The $p$-values and information on these SNPs are provided in Table S2. The $x$-axis contains the counts of the alternative allele, and the $y$-axis contains the counts of the reference allele. The dashed lines radiating from the origin are the expected counts under the fitted model of updog. The colors indicate posterior mode genotype. The "+" and "×" symbols indicate the parents.

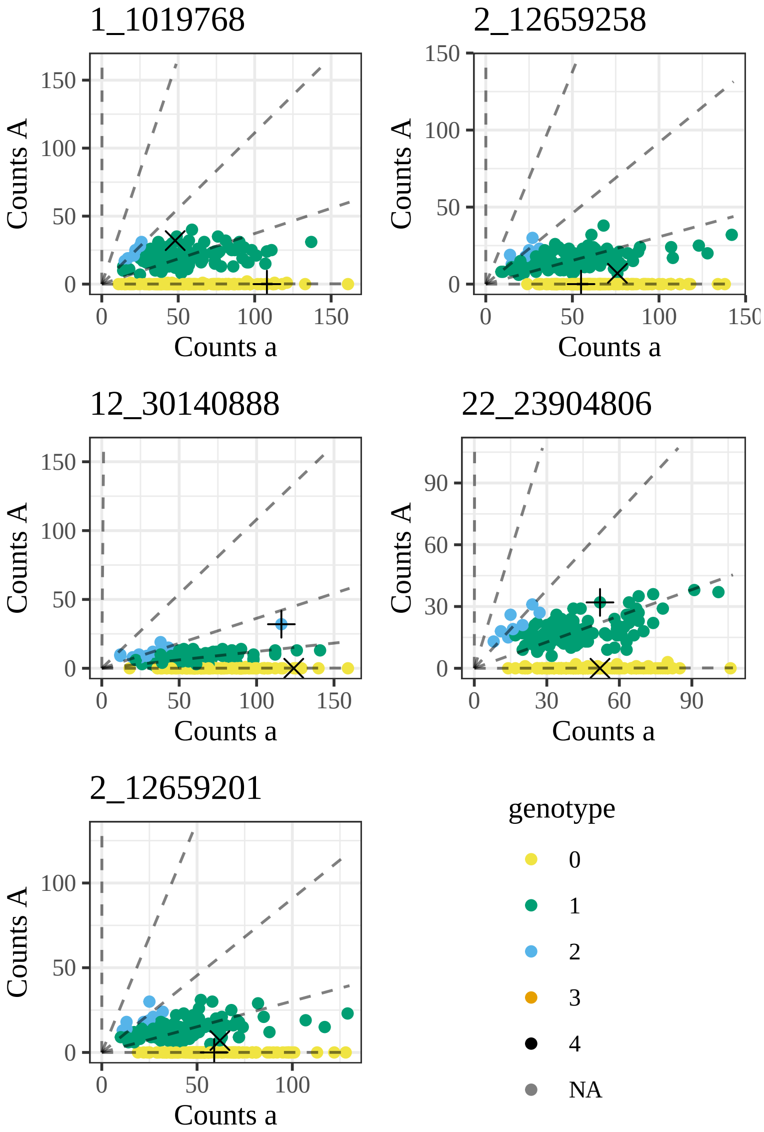


Figure S12: Genotype plots [Gerard et al., 2018] of five SNPs (titles) where the LRT indicates no segregation distortion, but `polymapR` indicates strong segregation distortion. The $p$-values and information on these SNPs are provided in Table S2. The $x$-axis contains the counts of the alternative allele, and the $y$-axis contains the counts of the reference allele. The dashed lines radiating from the origin are the expected counts under the fitted model of `updog`. The colors indicate posterior mode genotype. The "+" and "×" symbols indicate the parents.
